# Axonal defasciculation is restricted to specific branching points during regeneration of the lateral line nerve in zebrafish

**DOI:** 10.1101/2025.07.23.666336

**Authors:** Rohan S. Roy, A. J. Hudspeth

**Affiliations:** Laboratory of Sensory Neuroscience, The Rockefeller University, New York, NY 10065, USA; Howard Hughes Medical Institute, The Rockefeller University, New York, NY 10065, USA

**Keywords:** Hair cell, Collagen XVIII, Synapse, Axon guidance, Schwann cell, Extracellular matrix

## Abstract

Peripheral nerve regeneration requires precise selection of the appropriate targets of innervation, often in an environment that differs from that during the developmental wiring of the neural circuit. Severed axons of the zebrafish posterior lateral line nerve have the capacity to reinnervate mechanosensory hair cells clustered in neuromast organs. Regeneration represents a balance between fasciculated regrowth of the axonal bundle and defasciculation of individual axons into the epidermis where neuromasts reside. The cues that guide pathfinding during regeneration of the posterior lateral line nerve are unknown. Here we show that expression of *col18a1a,* which codes for the secreted heparan sulfate proteoglycan collagen XVIII, biases axonal defasciculation to specific branching points that coincide with circumscribed gaps in the epidermal boundary. We found that *col18a1a* is expressed by the neuromast and by a subset of Schwann cells that are located at the points of axonal defasciculation. Furthermore, we observed axon branching at inappropriate locations during nerve regeneration in *col18a1a* mutants. We propose a model in which a collagen XVIII-based axon-guidance cue complex attracts defasciculated axons across the epidermal basement membrane.

**Summary Statement:** The success of nerve regeneration depends on precise axon pathfinding and accurate target selection. We identify neuron-extrinsic factors that guide regeneration in a zebrafish model.

## Introduction

Damage to sensory neurons of the peripheral nervous system is often accompanied by detachment from end organs and degeneration of axon fragments distal to the site of injury (Waller, 1851). The ability of damaged peripheral axons to regenerate and reinnervate their original targets varies between vertebrates and depends on the site and type of injury (Deumens et al., 2010). In particular, regeneration of the peripheral neurites of spiral ganglion neurons in the mammalian cochlea is lacking. Retraction of afferent fibers from the sensory epithelium of the organ of Corti occurs directly through insult to spiral ganglion neurons or secondary to the death of the mechanosensitive hair cells that are presynaptic to afferent terminals (Liberman, 2017; Shibata et al., 2011). Regardless of the cause, the retraction of afferent endings is irreversible and results in permanent sensorineural hearing loss.

Compared to mammals, the zebrafish (*Danio rerio*) possesses a greater capacity for regeneration of both the central and peripheral nervous system (Rasmussen & Sagasti, 2017). The posterior lateral line (pLL) of the zebrafish has emerged as a model for studying the regeneration of peripheral sensory nerves (Ceci et al., 2014; Lozano-Ortega et al., 2018; Villegas et al., 2012; Xiao et al., 2015). The pLL comprises discrete neuromast organs that are deposited by a migratory group of primordial cells on the lateral surface of each side of the fish (Metcalfe, 1985). Each neuromast includes 10 to 20 sensory hair cells clustered in a rosette and surrounded by non-sensory supporting cells (Ghysen & Dambly-Chaudière, 2007). Neuromast hair cells, which are genetically, structurally, and functionally similar to those of the inner ear, are polarized to detect the direction of external water flow and thus to mediate swimming behaviors such as rheotaxis, schooling, and predator avoidance (Pickett & Raible, 2019).

Neuromasts are innervated by the pLL nerve, which contains predominantly afferent fibers postsynaptic to hair cells and a few efferent fibers presynaptic to hair cells (Haehnel et al., 2012; Haehnel-Taguchi et al., 2018; Manuel et al., 2021). On each side of a larva, the cell bodies of approximately 40 to 60 afferent neurons reside in a pLL ganglion (Haehnel et al., 2012; Raible & Kruse, 2000). These bipolar neurons have short central projections to the hindbrain and long peripheral axons that extend along the horizontal myoseptum as a nerve from which individual fibers defasciculate serially to innervate neuromasts (**Fig. 1A**) (Lopez-Schier & Pujol-Martí, 2013). Although peripheral axons are initially towed by a primordium lateral to the epidermal basement membrane, axon shafts are repositioned medial to the basement membrane whereas defasciculated arbors remain embedded in the neuromast (Raphael et al., 2010).

**Figure 1.**
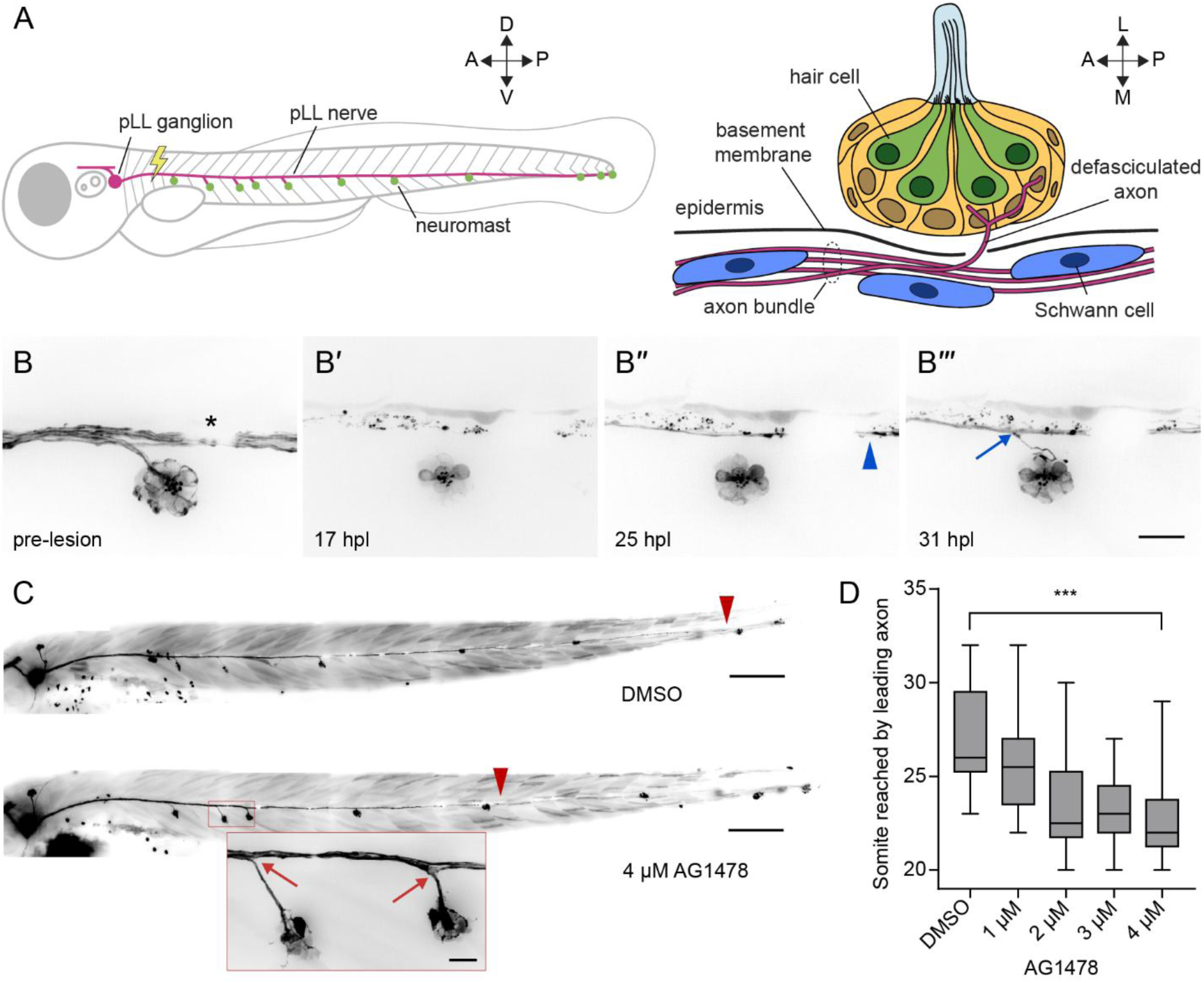
Regeneration of the pLL nerve occurs in stages. (A) A diagram of the larval zebrafish pLL (left, parasagittal view) and an individual neuromast (right, coronal transverse view). For all experiments, the nerve was lesioned at the level of the fourth somite. A – anterior, P – posterior, D – dorsal, V – ventral, L – lateral, M – medial. (B) Individual frames from a timelapse video portray the reinnervation of a neuromast following nerve lesion in a 4 dpf *Tg(HGn39d; myo6b:actb1-EGFP)* larva. (B’) Wallerian degeneration occurred within 24 hours post lesion (24 hpl) and was followed by passage of a leading axon (blue arrowhead) past the denervated hair cells of the neuromast (B’’). Six hours later, an individual follower axon defasciculated (blue arrow) from the axon bundle to enter the neuromast (B’’’). An asterisk marks a melanocyte that blocked underlying fluorescence. Scale bar, 20 µm. (C) Representative images show 6 dpf *Tg(HGn39d)* larvae two days after nerve lesion and recovery in either 1% DMSO (top) or 4 µM ErbB tyrosine kinase inhibitor AG1478 (bottom). Leader axons are marked with red arrowheads and defasciculation of follower axons marked with red arrows (inset). Scale bar, 100 µm, inset scale bar, 20 µm. (D) A box-and-whisker plot of the somite reached by the leading axon 2 dpl and recovery in different concentrations of AG1478. Whiskers span minimal and maximal values. *** *p*-value < 0.001, unpaired *t*-test. n = 16 zebrafish for DMSO, 1 µM, 2 µM, 3 µM, and 4 µM AG1478. n = 14 zebrafish for 1 µM AG1478.

The larval pLL offers a simple and accessible neural circuit in which to study axonal pathfinding, target selection, and hair cell reinnervation following nerve lesion. Regeneration of the larval nerve is quick: an axotomized nerve can reinnervate all neuromasts within two to three days post lesion (2-3 dpl) (Ceci et al., 2014; Villegas et al., 2012; Xiao et al., 2015). Concomitant with Wallerian degeneration of distal fragments, cut axons first bridge the lesion with the assistance of neighboring Schwann cells (Ceci et al., 2014). Following the retrograde clearance of axonal debris, the proximal stump regenerates along stripes of surviving Schwann cells, the bands of Büngner (Ceci et al., 2014; Xiao et al., 2015). Most cut axons reinnervate neuromasts distinct from those that they initially innervated during development, albeit with a preservation of global somatotopy (Ceci et al., 2014; Lozano-Ortega et al., 2018). This pattern suggests a competition between neuron-intrinsic and -extrinsic cues to the regenerating growth cone that affects neuromast selection. Only a few axonal growth cones defasciculate from the bundle at specific locations to enter the epidermis, pierce a basal layer of supporting cells, and reinnervate hair cells. The factors that bias growth-cone decisions at each of these steps remain unknown.

The extracellular matrix (ECM) also plays an instructive role in the guidance of axons through mechanical and chemical signaling. Secreted heparan sulfate proteoglycans (HSPGs), glycoproteins with covalently linked polysaccharide heparan sulfate side chains, are embedded in the ECM and exhibit a variety of cell-signaling properties. HSPGs can signal directly through domains on the core protein and moieties on its sugar side chains, or indirectly through secondarily bound ligands (Poulain & Yost, 2015; Sarrazin et al., 2011). Collagen XVIII is one such secreted HSPG that has been observed in epithelial and vascular basement membranes throughout the body as well as in the peripheral nervous system (Halfter et al., 1998; Seppinen & Pihlajaniemi, 2011). In zebrafish, the gene *col18a1*, which codes for the α1 chains of collagen XVIII, is needed for the ventral extension of motor growth cones from the spinal cord into the musculature and for the regeneration of severed retinal ganglion cell axons across the optic chiasm (Harvey et al., 2024; Schneider & Granato, 2006). Collagen XVIII and its homologues have also been shown to affect axonal guidance in *Caenorhabditis elegans* (Ackley et al., 2001) and *Drosopholia melanogaster* (Meyer & Moussian, 2009), as well as in the enteric nervous systems of the chick and mouse (Nagy et al., 2018). These observations suggest a conserved role in the patterning of the nervous system.

In the present study, we used a combination of timelapse imaging, bulk RNA sequencing, and RNA fluorescence *in situ* hybridization (RNA-FISH) to characterize the defasciculation of individual axons from the regenerating axon bundle into the denervated neuromast. We find that *col18a1a* is expressed by the neuromast and a subset of Schwann cells that lie at the point of axon defasciculation. Using a transgenic line that labels the epidermal boundary, we show that axon defasciculation occurs through specific gaps in the ECM. We additionally generated a *col18a1a* mutant that exhibits axonal branching at inappropriate locations during nerve regeneration. We propose a model in which extracellular HSPG-axon guidance cue complexes form a signaling pathway to restrict axonal defasciculation toward neuromast targets in the epidermis.

## Results

### Regeneration of the pLL nerve occurs in stages

Peripheral afferent axons of the larval pLL nerve extend from somata in the pLL ganglion to the caudal end of the fish, innervating seven to eleven neuromasts along the horizontal myoseptum. Individual axons defasciculate from the nerve to enter the epidermis and innervate hair cells located in the center of each neuromast (**Fig. 1A**). To label lateral line afferent axons and hair cell membranes for timelapse imaging, we used the *Tg(HGn39d; myo6b:actb1-EGFP)* transgenic line (Kindt et al., 2012; Nagayoshi et al., 2008). In each larva four days post fertilization (4 dpf), we lesioned a 20 µm section of the pLL nerve between the ganglion and the first neuromast with a 355 nm laser. We then imaged the reinnervation of hair cells in the neuromast posterior to the cut site (**Fig. 1B**). As previously shown (Ceci et al., 2014), axon severing was followed by Wallerian degeneration of the axonal fragments distal to the cut site. As debris was being cleared in the 24 hours following nerve lesion, a few pioneering axons extended past the denervated neuromast. About six hours after the first axons bypassed the neuromast, a follower axon defasciculated ventrolaterally from the regenerating axon bundle to reinnervate the hair cells of the neuromast.

To further distinguish the stages of pLL nerve regeneration, we perturbed regeneration by administering the ErbB tyrosine kinase inhibitor AG1478 following nerve lesion. Early administration of AG1478 blocks Schwann cell migration along the peripheral axons of the pLL nerve through interference of ErbB-neuregulin signaling (Lush & Piotrowski, 2014; Lyons et al., 2005). AG1478 administered after Schwann cells had migrated along the entirety of the nerve delayed axon traversal of the cut site following axotomy, but not the velocity of axon growth (Ceci et al., 2014). We asked whether blocking ErbB signaling could affect other stages of regeneration, especially the defasciculation of individual axons toward the neuromast. Recapitulating earlier results, we observed a significant decrease in the distance along the trunk of the fish that the leader axon had reached by 2 dpl (**Fig. 1 C, D**). Despite the delay to leader axon growth, we observed normal bundled growth along the horizontal myoseptum and defasciculation of follower axons from the bundle to reinnervate neuromasts (**Fig. 1C**). Although ErbB signaling might be involved in the initial passage of severed axons across the lesion site, it is dispensable for neuromast reinnervation. Our live imaging indicated that nerve regeneration proceeds in two stages following axon traversal of the lesion site: the extension of the axon bundle along the horizontal myoseptum in a leader-follower fashion and the defasciculation of follower axons from the bundle at specific locations along the trunk. The different locations and timing of these stages suggest that they are guided by distinct signaling cues.

### Neuromast hair cells transiently change gene expression after denervation

To identify guidance cues that may be coming from the targets of innervation, we performed bulk RNA sequencing on denervated hair cells. At either 1 dpl or 3 dpl, hair cells from *Tg(neurod:tdTomato; myo6b:actb1-EGFP)* transgenic larvae were isolated and sequenced; the results were compared to those from innervated hair cells of age-matched larvae in which a sham lesion had been performed (**Fig. 2A**). Prior to fluorescence-activated cell sorting (FACS), larvae were decapitated to exclude hair cells from the otic vesicle, anterior lateral line, and dorsal branch of the pLL. At 1 dpl, leader axons had only begun to extend along the horizontal myoseptum and most neuromast hair cells were denervated (**Supplemental Fig. 1**). By 3 dpl, the majority of neuromast hair cells had been reinnervated and there was functional recovery of swimming behavior. Hair cells were sorted based on size and GFP expression (**Supplemental Fig. 2A**), and their identity was confirmed *post hoc* in the bulk sequencing data by cross-referencing the most-expressed genes to a published single-cell RNAseq dataset of the neuromast (**Supplemental Fig. 2B-C**) (Baek et al., 2022).

**Figure 2.**
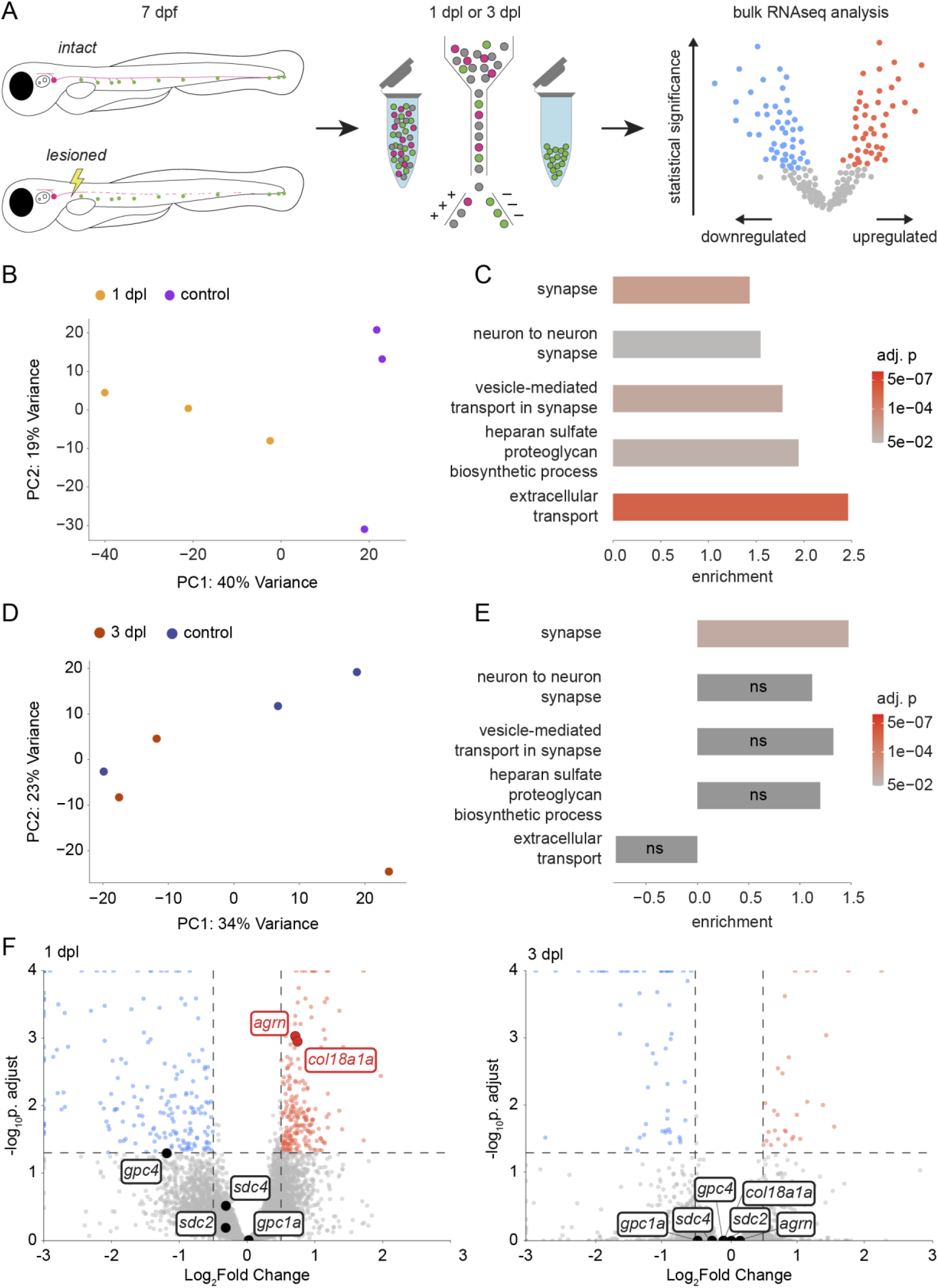
Neuromast hair cells transiently change gene expression upon denervation. (A) A schematic diagram of the hair cell-sequencing workflow. The pLL nerve of *Tg(neurod:tdTomato; myo6b:actb1-EGFP)* larvae was either lesioned or left intact at 7 dpf. Hair cells were isolated either 1 dpl or 3 dpl and pooled together for bulk sequencing. (B) A principal-component analysis plot of samples from denervated hair cells sequenced 1 dpl (n = 4,058 cells, 4,833 cells, and 3,723 cells) and control hair cells sequenced 1 day post sham lesion (n = 5,973 cells; 6,045 cells; and 2,744 cells). (C) Gene set enrichment analysis of select synapse (GO:0045202, GO:0098984), extracellular transport (GO:0099003, GO:0006858), and HSPG synthesis (GO:0015012) gene sets in denervated hair cells 1 dpl compared to age-matched controls. (D) A principal component analysis plot of samples from hair cells sequenced 3 dpl (n = 1,735 cells; 4,363 cells; and 3,901 cells) and control hair cells sequenced 3 days post sham lesion (n = 5,123 cells; 2,482 cells; and 4,861 cells). (E) Gene set enrichment analysis of the same GO sets in (C) in hair cells 3 dpl compared to age-matched controls. ns – not significant, adjusted *p-value* > 0.05. (F) A volcano plot of differential gene expression in hair cells 1 dpl (left) or 3 dpl (right) compared to age-matched controls (red – upregulated, blue – downregulated, gray – not significant), with genes coding for transmembrane and secreted HSPG core proteins highlighted. Statistical significance set at a *p-value* of 0.05 adjusted for multiple comparisons (horizontal dashed line) and biological significance set at 0.5 log2-fold change (vertical dashed line).

Upon dimensionality reduction of the data through principal component analysis, hair cell samples from larvae 1 dpl clustered separately from control samples along the first principal component axis, whereas hair cell samples from larvae 3 dpl were intermixed with those from controls (**Fig. 2B, D**). Furthermore, there were more genes differentially expressed in hair cells 1 dpl compared to 3 dpl (**Supplemental Fig. 3A, B**). These summary results imply that there are significant changes in the transcriptome of hair cells in response to denervation that revert after hair cells have been reinnervated.

We observed significant enrichment of synapse and extracellular transport gene-ontogeny sets (GO:0045202, GO:0098984, GO:0099003, GO:0006858) in denervated hair cells 1 dpl that, with the exception of the synapse gene set (GO:0045202), was lost by 3 dpl (**Fig. 2C, E**). There was no significant upregulation of genes coding for canonical axon guidance cues in the denervated hair cells, but *semaphorin7a (sema7a)*, *brain-derived neurotrophic factor (BDNF)*, and *reticulon4a (rtn4a)* were highly expressed across all hair cell samples (**Supplemental Fig. 4**). The gene set for the biosynthesis of heparan sulfate proteoglycans (GO:0015012), which comprises genes for sulfotransferases, glycosyltransferases, and other enzymes that modify heparan sulfate side chains, was transiently enriched in denervated hair cells (**Fig. 2C, E**). There was additionally a transient upregulation of the genes coding for the secreted HSPG core proteins agrin (*agrn*) and collagen XVIII (*col18a1a*), but not of the transmembrane HSPG core proteins syndecan and glypican (**Fig. 2F**).

Our bulk sequencing data indicate that a transient change in the gene expression of denervated hair cells is in part due to recuperating lost synapses. The upregulation of secreted HSPGs and their modifying enzymes suggests that these cell-signaling molecules influence pLL axonal guidance. Although *agrn* is expressed throughout hair cells and supporting cells of the neuromast, we did not observe pathfinding defects by afferent axons in *agrn^p168^* mutants following nerve lesion (Gribble et al., 2018) (**Supplemental Fig. 5**). We accordingly focused on characterizing the role of *col18a1a* during nerve regeneration.

### *col18a1a* is expressed in the neuromast and in Schwann cells at the points of axonal defasciculation

To further investigate the timing and location of *col18a1a* expression, we performed RNA-FISH on innervated and denervated neuromasts. We observed diffuse expression of *col18a1a* RNA throughout the innervated neuromast, both in hair cells and in supporting cells (**Fig. 3A**). We also noted strong expression of *col18a1a* in a *sox10^+^* Schwann cell positioned outside the neuromast at the location where an individual axon defasciculated from the nerve. Although *sox10^+^* Schwann cells lined the entirety of the pLL nerve, we counted only a single Schwann cell expressing *col18a1a* at the point of axonal defasciculation in 40 of 46 neuromasts imaged (**Fig. 3D, E**).

**Figure 3.**
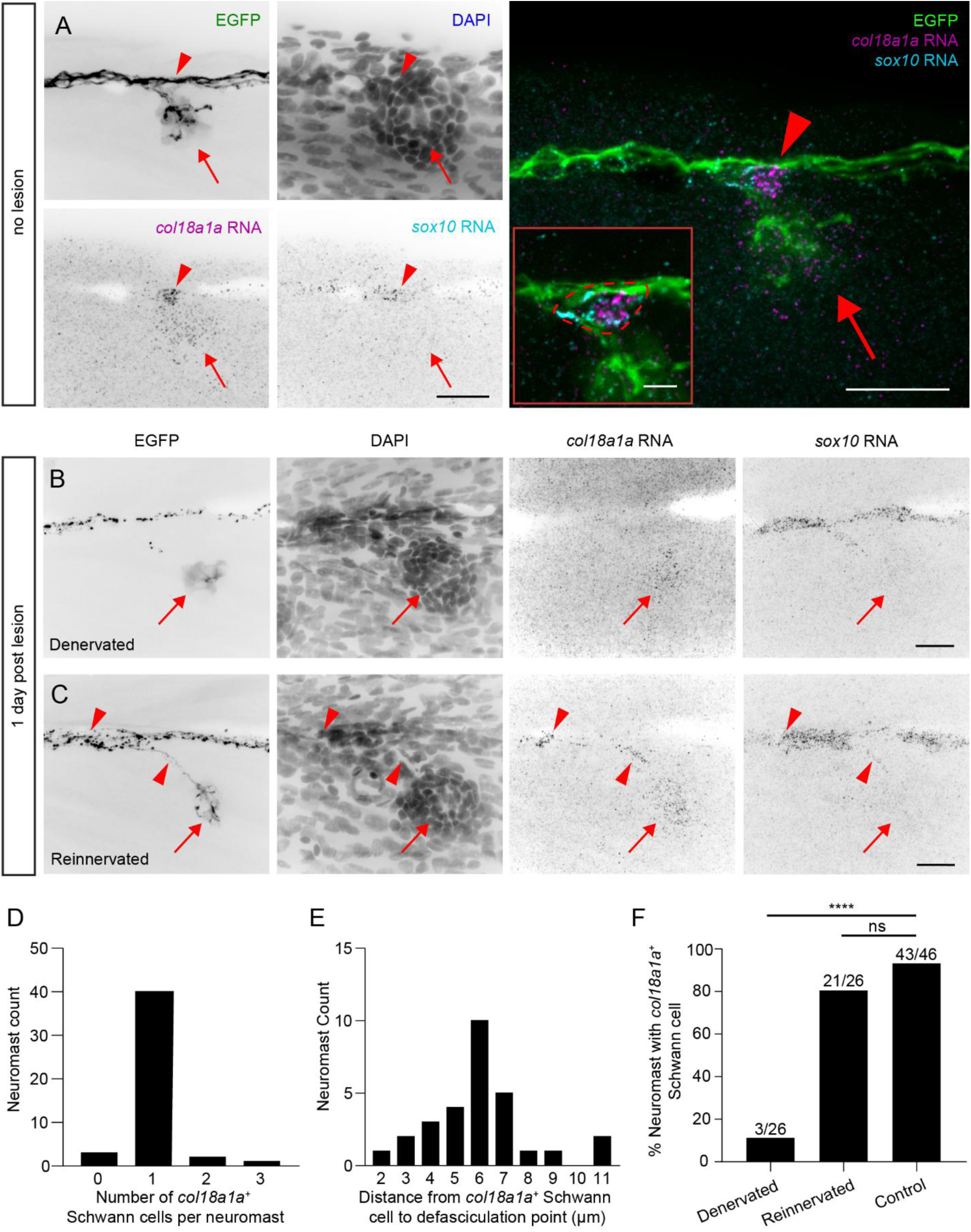
Spatiotemporal expression of *col18a1a* RNA in neuromasts and a subset of Schwann cells. (A) A maximum projection of individual color channels (left) and a merged image (right) of RNA-FISH against *col18a1a* and *sox10* in a *Tg(HGn39d; myo6b:actb1-EGFP)* larva fixed 8 dpf. *col18a1a* is expressed throughout the neuromast (red arrow) and in a single Schwann cell (red arrowhead). Inset – A single imaging plane of the *col18a1a^+^/sox10^+^* Schwann cell (dotted red outline). Scale bar of individual color channels and merged image, 20 µm. Inset scale bar, 5 µm (B) A denervated neuromast 1 dpl. There is expression of *col18a1a* only in the neuromast (red arrow). Scale bar, 20 µm. (C) A reinnervated neuromast from the larva shown in panel (B). *col18a1a* is expressed in the neuromast (red arrow) and in two Schwann cells adjacent to axon branching (red arrowheads). Scale bar, 20 µm. (D) The number of *col18a1a^+^* Schwann cells per neuromast in 8 dpf larvae. n = 46 neuromasts from 27 larvae. (E) The distance from the point of axon defasciculation to the nearest *col18a1a^+^* Schwann cell. n = 29 primary, non-terminal neuromasts from 21 larvae. (F) The proportion of denervated, reinnervated, and control neuromasts with at least one adjacent *col18a1a^+^* Schwann cell. n = 26 denervated neuromasts from 19 larvae, 26 reinnervated neuromasts from 19 larvae, and 46 control neuromasts from 27 larvae. **** *p-value* < 0.0001 by Fisher’s exact test. ns, not significant.

We probed neuromasts at 1 dpl to assess any changes in the spatiotemporal expression of *col18a1a* during pLL nerve regeneration. We were able to document *col18a1a* expression in denervated and recently reinnervated neuromasts within the same larva. We detected the same spatial pattern of *col18a1a* expression in denervated neuromasts and reinnervated neuromasts that we observed in control neuromasts. The high variability of probe intensity and background signal precluded our quantification of differences in fluorescence between neuromasts. However, we did note an absence of *col18a1a* expression in Schwann cells adjacent to denervated neuromasts (**Fig. 3B, F**). In the same fish, in a neuromast that had been reinnervated, we again detected the presence of at least one *col18a1a^+^* Schwann cell at the axon branch point (**Fig. 3C, F**). Based on these results, we reasoned that dynamic expression of *col18a1a* by specialized Schwann cells at specific choice points biases axon defasciculation towards the neuromast.

To further investigate the relationship between pLL axons and the ECM at branching points, we used a *Tg(krt19:col1α2-GFP; hsp70:NTR2.0-2A-mCherry en.Sill1)* transgenic line, henceforth referred to as *Tg(krt19:col1α2-GFP, sill1:mCherry)*. This line expresses the fluorescent protein mCherry in lateral line afferents (Pujol-Marti et al., 2012) and a secreted collagen I-GFP fusion protein that incorporates into a collagen I mesh directly underneath the epidermal basement membrane (Morris et al., 2018). At 5 dpf, we observed well-circumscribed gaps in the collagen I mesh through which defasciculated axons passed to innervate hair cells of primary neuromasts residing in the epidermis (**Fig. 4A**). Each gap had an area of 13.24 ± 5.13 µm^2^ and was positioned close to the nerve, within 4.24 ± 1.95 µm of axon defasciculation (mean ± SD, **Fig. 4B, C**). In secondary neuromasts, which are deposited by a second migrating primordium and lie farther ventral from the nerve compared to primary neuromasts, defasciculated axons travelled within the collagen I mesh, rather than through a circumscribed gap (**Supplemental Fig. 6A-C**). Although the entrance to the collagen I mesh lay where axons defasciculated from the nerve and the exit corresponded to the neuromast’s location in the epidermis, these locations were variable and difficult to locate precisely. For this reason, we restricted further analysis to primary neuromasts, for which a circumscribed gap could always be clearly defined.

**Figure 4.**
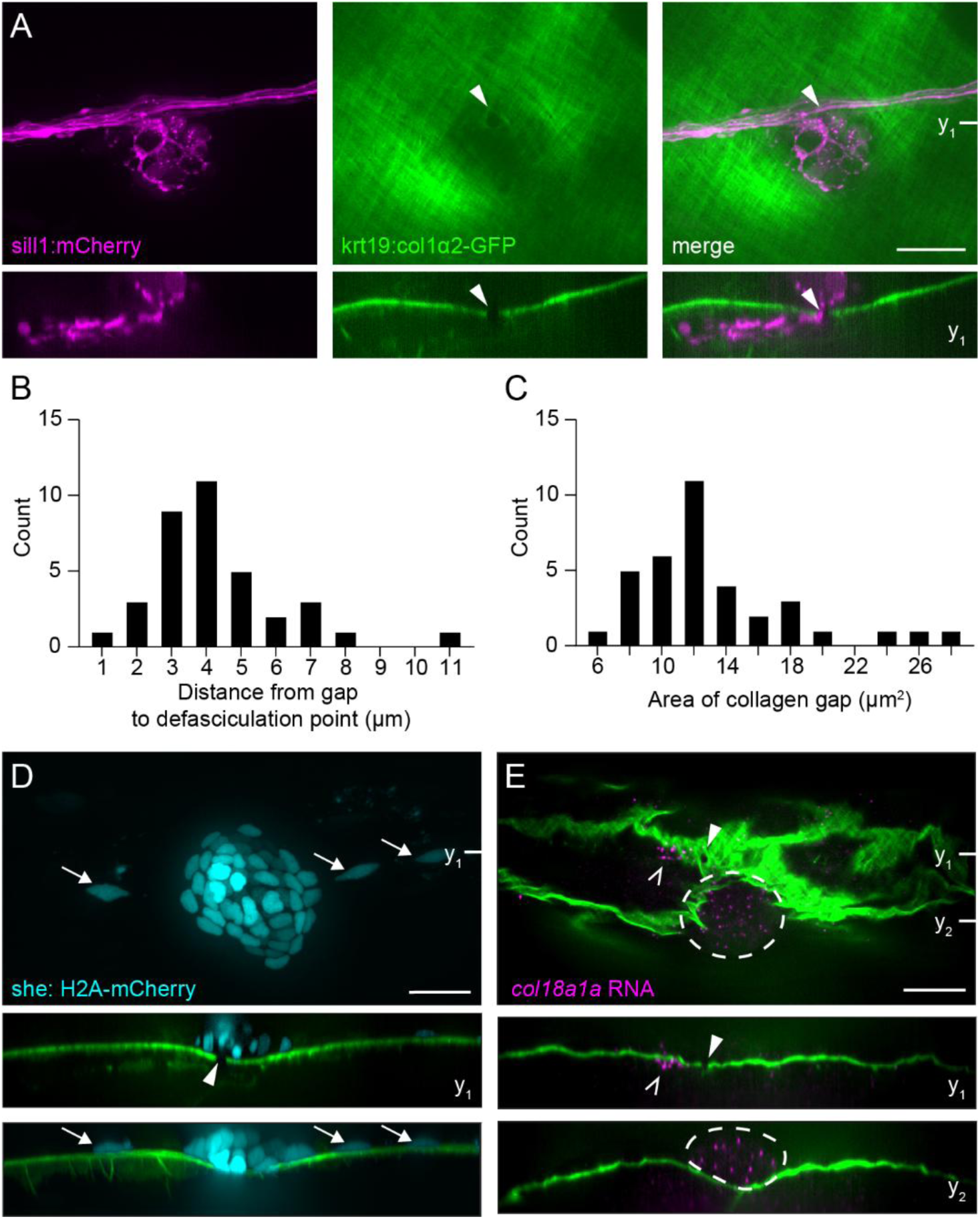
Axons cross gaps in the epidermal boundary layer to innervate neuromast. (A) A parasagittal projection (top) and a coronal slice (bottom) taken at the indicated plane of a confocal stack imaged in a live 5 dpf *Tg(krt19:col1α2-GFP, sill1:mCherry)* larva. Individual afferents labeled by the *Sill1* enhancer (magenta) defasciculate from the nerve and pass through a gap (white arrowhead) in the collagen I matrix (green). (B) The distribution of distances between the ECM gap and point of axon defasciculation in 5 dpf larvae. n = 36 zebrafish. (C) The distribution of gap areas. n = 36 zebrafish. (D) A sagittal projection (top), coronal slice (middle) taken at the indicated plane, and a coronal projection (bottom) of a confocal stack imaged in a live 4 dpf *Tg(krt19:col1α2-GFP, she:H2A-mCherry)* larva. Neuromast (cyan) and adjacent interneuromast cells (cyan, white arrows) reside in the epidermis, lateral to the collagen I matrix and gap. mCherry fluorescence intensity gamma corrected. (E) An individual sagittal slice (top) and coronal slices (middle, bottom) taken at the planes indicated of RNA-FISH against *col18a1a* in a fixed, 8 dpf *Tg(krt19:col1α2-GFP)* larva. A single *col18a1a^+^* cell resides medial to the epidermal boundary layer (white caret) and adjacent to the ECM gap (white arrowhead). Cells of the neuromast (white dotted outline) are also *col18a1a^+^* and reside lateral to the epidermal boundary. All scale bars, 20 µm.

In contrast to the axon bundle, which coursed medial to the epidermis (**Fig. 4A**), neuromast cells and adjacent interneuromast cells labeled in *Tg(she: H2A-mCherry, krt19:col1α2-GFP)* fish were positioned lateral to the basement membrane (**Fig. 4D**). To localize *col18a1a* expression in relation to the ECM, we performed RNA-FISH on fixed 8 dpf *Tg(krt19:col1α2-GFP)* larvae. Although the fixation and *in situ* protocol distorted the ECM, we could nonetheless observe a single *col18a1a*^+^ cell directly medial to the collagen I mesh and adjacent to a gap. In comparison, the *col18a1a* expression in the neuromast occurs lateral to the basement membrane (**Fig. 4E**). The constrained expression of *col18a1a* across the epidermal boundary layer, in proximity to the gaps through which defasciculated axons pass, further indicates a role in axonal guidance.

### Branching axons of the regenerating pLL nerve extend through pre-formed gaps in the extracellular matrix

The gaps that permit extension of defasciculated axons into the epidermis are a consequence of basement membrane remodeling that occurs during development (Raphael et al., 2010). We further investigated the role of these gaps during nerve regeneration. We imaged the innervation of primary neuromasts through these gaps in 5 dpf *Tg(krt19:col1α2-GFP; sill1:mCherry)* larvae, lesioned the nerve, and reimaged the same gaps 1 dpl and 3 dpl (**Fig. 5A’-A’’**). In 31 of 32 larvae, we observed regeneration through the same ECM gap through which the axons extended before lesioning. In the one exceptional case, axons traversed a different preexisting gap in the ECM. Gaps remained the same size and shape during the three-day course of nerve regeneration, and we did not observe the formation of any new gaps in this interval.

**Figure 5.**
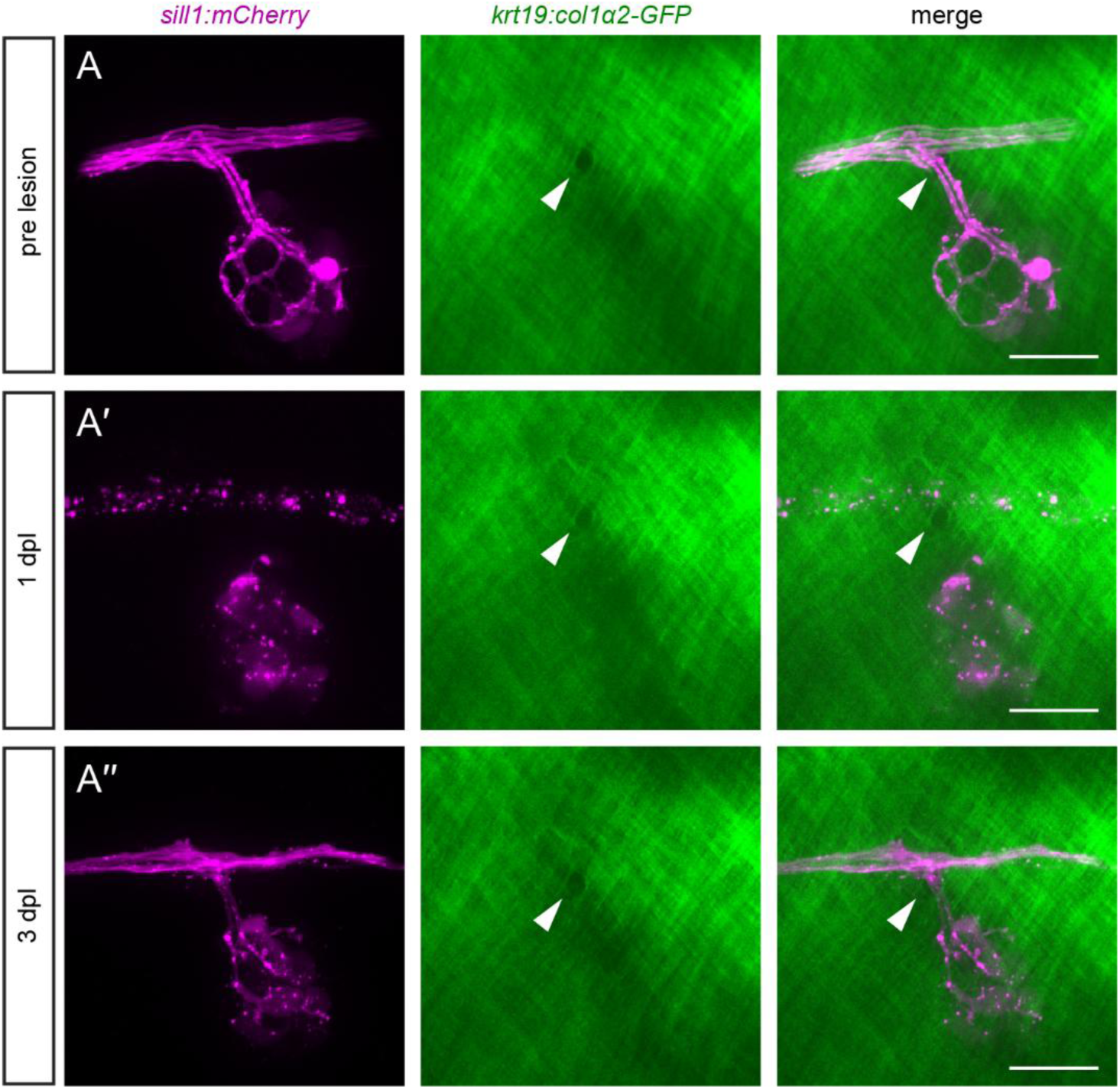
Defasciculation through a boundary gap during axon regeneration. (A) Before nerve lesion, a primary neuromast in a 5 dpf *Tg(krt19:col1α2-GFP, sill1:mCherry)* larva was innervated through a gap in the ECM (white arrowhead). (A’) While axons underwent Wallerian degeneration at 1 dpl, the gap in the ECM remained unchanged. (A’’) By 3 dpl, axons had reinnervated the neuromast by defasciculating through the original ECM gap. Scale bar, 20 µm.

To determine whether axons would extend into the epidermis through any gap in the ECM, we punctured the epidermal boundary layer with a glass microneedle somewhere along the path of the regenerating nerve. Each puncture created an approximately 30 µm-wide gap in the ECM. There was full extension of an axon into the epidermis through this artificial gap in only 1 of 33 larvae imaged. For the other 32 larvae, the majority of axons remained tightly fasciculated beneath the epidermal boundary, with the occasional axon stalled along the edges of the gap (**Fig. 6A**).

**Figure 6.**
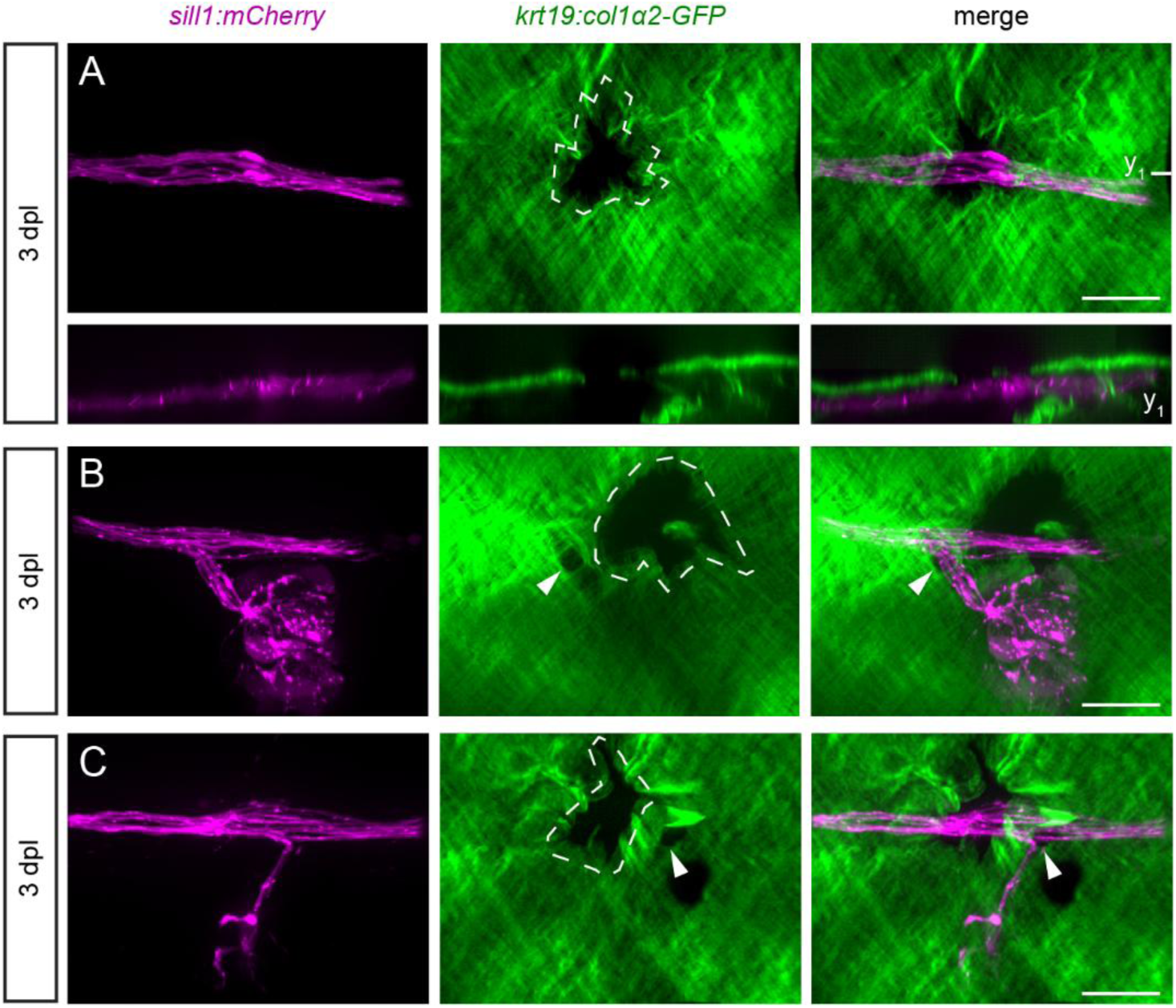
Axons do not defasciculate through artificial gaps in the ECM. (A) A parasagittal projection (top) and coronal slice at the plane indicated (bottom) of a regenerated nerve bypassing a puncture made in the ECM (dotted white outline) in an 8 dpf *Tg(krt19:col1α2-GFP, sill1:mCherry)* larva. Scale bar, 20 µm. (B) In another larva, the reinnervation of a neuromast proceeded through a natural ECM gap (white arrowhead) rather than through a puncture made just posterior to the gap. Scale bar, 20 µm. (C) Reinnervation of a neuromast in a different larva similarly bypassed a puncture made just anterior to the natural ECM gap. Scale bar, 20 µm.

In some larvae, the puncture was made along the path of the regenerating nerve, within 50 µm of a neuromast. In these cases, we could test whether the regenerating axons would pass through the original gap or the artificial gap to reinnervate the neuromast. Because of their size and irregular borders, the artificial gaps were easy to distinguish from the smaller, more circular gaps that naturally formed during development. We found that axons preferred to reinnervate neuromasts through the natural gaps compared to the artificial gaps (10 out of 11 larvae). This was true whether the artificial gap was located posterior (five of five larvae, **Fig. 6B**) or anterior (five of six larvae, **Fig. 6C**) to the natural gap. The preference of an axon to defasciculate through the natural gap in the ECM suggests that there is an attractive signal specifically at this location that draws growth cones into the epidermis.

### *col18a1a* mutants have aberrant axon pathfinding during pLL nerve regeneration

Based on the spatiotemporal expression pattern of *col18a1a*, we hypothesized that this extracellular HSPG influences the defasciculation of individual axons into the epidermis. We generated a stable *col18a1a* mutant line by CRISPR/Cas9 gene editing (**Supplemental Fig. 7 A-C**). We selected mutants with a two-base-pair deletion in an exon common to all three isoforms of *col18a1a* and bred them into a *Tg(HGn39d; myo6b:actb1-EGFP)* background so as to have an afferent axon and a hair cell marker. Although innervation of neuromasts appeared grossly normal in homozygous mutants, the nerve displayed higher tortuosity compared to those of siblings (**Fig. 7 A, B**). Moreover, the nerve was less tightly packed together in homozygous mutants compared to those of siblings (**Fig. 7 F, G**).

**Figure 7.**
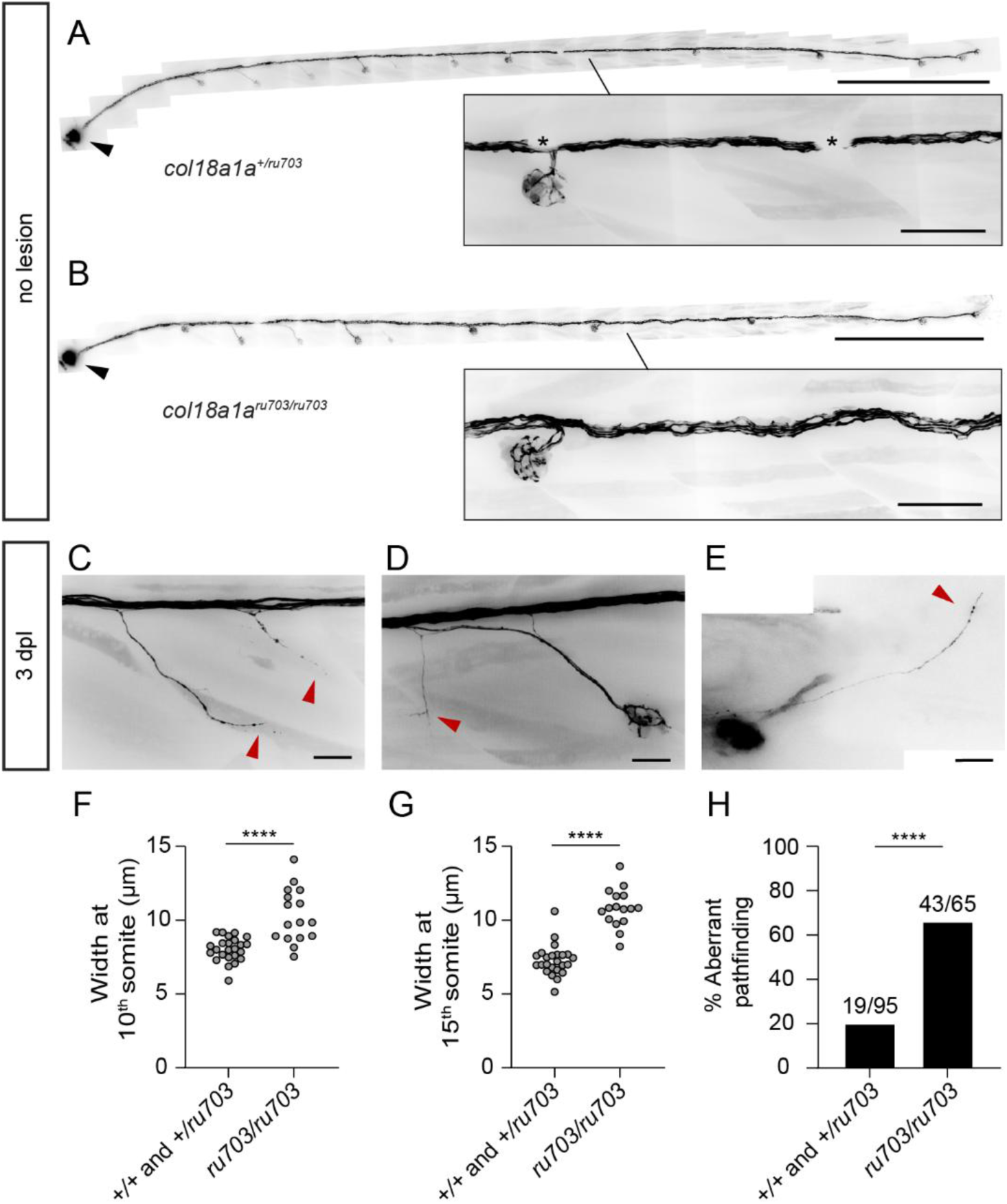
*col18a1a* mutants have aberrant axon pathfinding during pLL nerve regeneration. (A) The peripheral segment of the pLL nerve in an 8 dpf *col18a1a^+/ru703^ Tg(HGn39d; myo6b:actb1-EGFP)* larva. Asterisks denote melanocytes overlying the nerve. The pLL ganglion is marked by a black arrowhead in (A) and (B). Scale bar, 500 µm; inset scale bar, 50 µm. (B) In the peripheral segment of the pLL nerve in an 8 dpf *col18a1a^ru703/ru703^ Tg(HGn39d; myo6b:actb1-EGFP)* larva, the nerve is less tightly packed and follows a more tortuous path (inset). Scale bar, 500 µm; inset scale bar, 50 µm. (C) Aberrant defasciculation of axons (red arrowheads) in a mutant larva following nerve regeneration. Scale bar, 20 µm. (D) Normal reinnervation of a secondary neuromast accompanied with inappropriate collateral branching (red arrowhead). Scale bar, 20 µm. (E) Extension of a lone axon (red arrowhead) past the terminal neuromast. (F) The width of the axon bundle in 5 dpf *col18a1a^ru703/ru703^* mutants compared to control siblings at the level of the 10^th^ somite. **** *p-value*<0.0001, unpaired t-test. (G) The width of the axon bundle in 5 dpf *col18a1a^ru703/ru703^* mutants compared to control siblings at the level of the 15^th^ somite. **** *p-value<*0.0001, unpaired t-test. (H) The percentage of *col18a1a^ru703/ru703^* mutant larvae with aberrant axon pathfinding 3 dpl compared to control siblings. n = 95 wildtype/heterozygous larvae, 65 mutant larvae. **** *p-value*<0.0001, Chi-squared test.

We checked for abnormalities in *col18a1a^ru703/ru703^* mutants following pLL nerve lesion and axon regeneration. Although there was reinnervation of all neuromasts, we observed instances of individual axons defasciculating from the regenerated axon bundle at inappropriate locations. These axons would branch from the nerve at locations where there was no nearby neuromast (**Fig. 7C**). While we found examples of inappropriate defasciculation at different locations along the trunk, we noted that many abnormal branches surrounded the locations of secondary neuromasts (**Fig. 7D**). We also observed mutant larvae in which a lone axon extended past the terminal neuromast, into the developing tailfin (**Fig. 7E**). We counted significantly more mutant larvae with aberrant axon pathfinding compared to control siblings (**Fig. 7H**). Our results indicate that although *col18a1a* is not required for pLL nerve regeneration or hair cell reinnervation, it plays an auxiliary role in restricting axon defasciculation and subsequent pathfinding to appropriate locations.

## Discussion

Although the path that severed axons follow resembles the path established during development, there are key differences in how the growth cones reach their targets. The innervation pattern of neuromasts deposited during development reflects the chronology of neuron differentiation in the ganglion. Peripheral axons of early-differentiating neurons contact the primordium lateral to the epidermal basement membrane and project to the caudal end of the fish, whereas axons of later-differentiating neurons trail behind the primordium and innervate more rostral neuromasts (Pujol-Martí et al., 2010; Sato & Takeda, 2013). In contrast, during regeneration, axons must advance medial to the epidermal basement membrane in the absence of a primordium and instead extend along bands of Büngner (Xiao et al., 2015). Pioneering axons bypass denervated neuromasts to reach the caudal end of the fish (**Fig. 1B**). Follower axons have a simpler task of pathfinding by growth along the shaft of a pioneer. There is great plasticity in the rewiring of neuromasts following nerve lesion, with individual axons able to reinnervate different neuromasts from those that they originally innervated (Ceci et al., 2014; Lozano-Ortega et al., 2018). This pattern implies that the decision to defasciculate from the axon bundle is not hard-wired into an axon and is biased by external signaling cues.

To identify these cues, we first sequenced denervated hair cells in bulk to determine if there were transcriptional changes associated with axon guidance. We showed that hair cells transiently change their gene expression profile in response to denervation that reverts following reinnervation. Although we did not see an upregulation of canonical axon guidance cues, we did observe an upregulation of *col18a1a*, which codes for a secreted HSPG. HSPGs have an extensive role in neuronal migration, axon guidance, and synaptogenesis during nervous system development (Poulain & Yost, 2015). Once secreted from the cell, HSPGs alter the structure of basement membranes and provide storage depots for growth factors and other diffusible cues (Sarrazin et al., 2011). Although we saw additional branching at inappropriate locations during pLL nerve regeneration in our *col18a1a* mutants, we nevertheless observed proper axon defasciculation toward neuromasts. This behavior suggests an accessory role for collagen XVIII in refining axon branching rather than a primary guiding role.

We were surprised to find one to three *col18a1a^+^* Schwann cells lying outside a neuromast, at the location of axon defasciculation. Schwann cells have a multifaceted role in the regeneration of the pLL nerve. As in mammalian systems (Arthur-Farraj et al., 2012), zebrafish Schwann cells distal to the nerve lesion site de-differentiate and assume a pro-repair phenotype (Ceci et al., 2014; Xiao et al., 2015). Although not strictly necessary for axon extension, Schwann cells are required for the bundled growth of axons along the horizontal myoseptum (Ceci et al., 2014; Villegas et al., 2012). Our results suggest an additional role for *col18a1a^+^* Schwann cells that restrict the defasciculation of axons to specific choice points along the trunk of the fish. These choice points correspond to gaps in the ECM that allow passage of defasciculated axons into the epidermis. We propose that collagen XVIII, secreted by the neuromast and by specialized Schwann cells, scaffolds with axon guidance cues to create a signaling pathway from one side of the epidermal basement membrane to the other (**Fig. 8**). Potential candidates for complexed guidance cues include GDNF, a neurotrophic factor necessary for the association of towed axons with the primordium (Schuster et al., 2010), semaphorin 7a, a hair cell-derived chemoattractant (Dasgupta et al., 2024), BDNF, a neurotrophin expressed by hair cells and crucial for maintaining afferent innervation (Aragona et al., 2022; Mo & Nicolson, 2011), and reticulon 4/nogo-A, which has permissive axon guidance activity in zebrafish unlike its inhibitory effect in mammals (Abdesselem et al., 2009; Welte et al., 2015).

**Figure 8.**
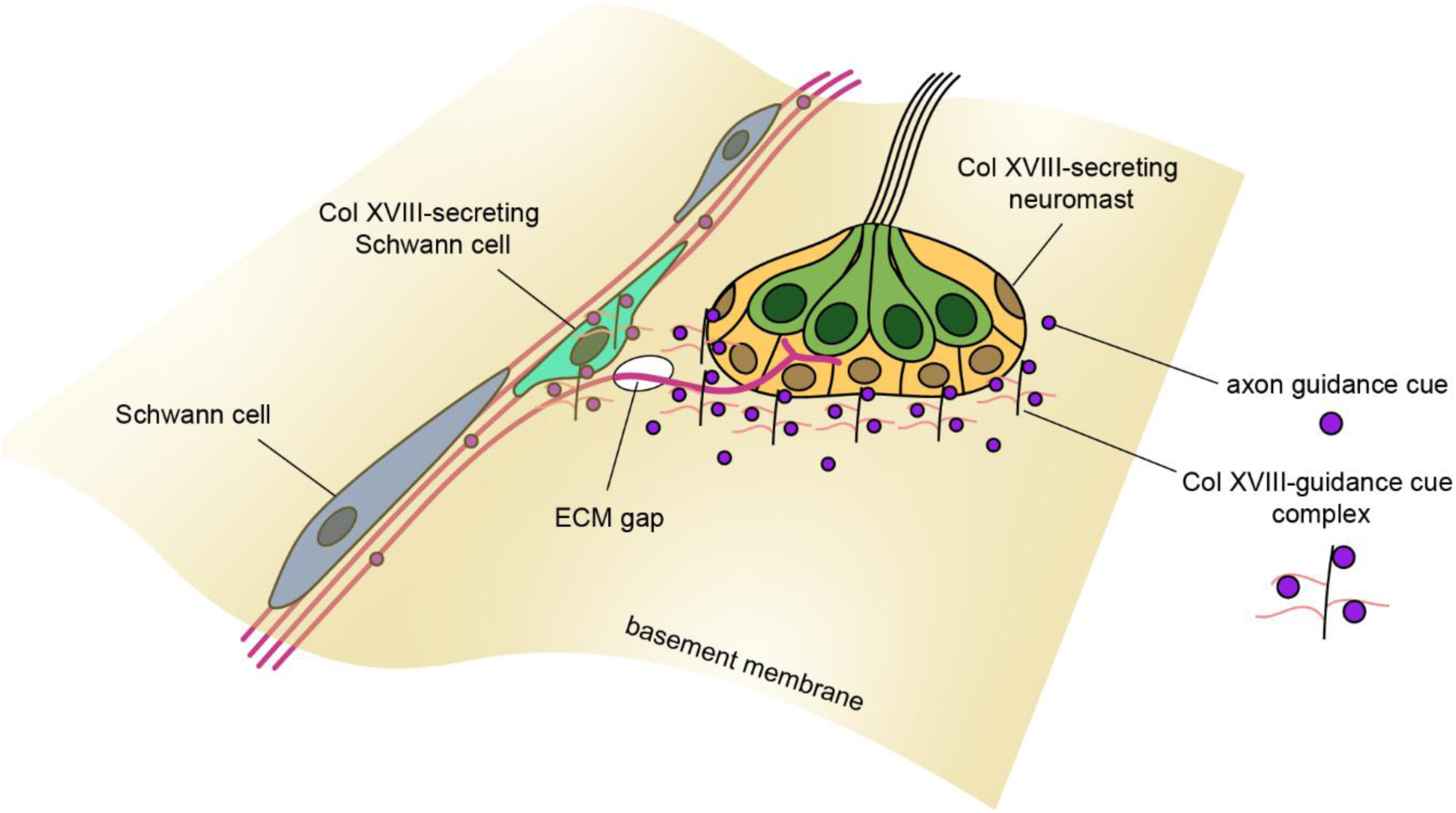
Proposed model of axon defasciculation. Collagen XVIII may act as an auxiliary scaffold for axon guidance cues to restrict defasciculation across the epidermal basement membrane to specific locations along the zebrafish trunk. Axon guidance cues could be complexed directly to the Collagen XVIII core protein or to attached heparan sulfate side chains.

Crossing the basement membrane represents a unique challenge to regenerating pLL axons. During development, the ECM is actively remodeled by Schwann cells to position the axon bundle beneath the epidermis and thereby isolates the nerve from postembryonic ventral migration of neuromasts (Raphael et al., 2010). This remodeling leaves small gaps in the ECM near the nerve through which defasciculated neurites enter the epidermis. Although these gaps can be seen in serial electron micrographs of the neuromast (Dow et al., 2018; Odstrcil et al., 2022; Raphael et al., 2010), there have been no studies visualizing these gaps in a live zebrafish. Using a transgenic line that expresses a collagen I-GFP fusion protein that incorporates into the epidermal boundary, we were able to visualize ECM gaps over the course of nerve regeneration. In contrast to the active remodeling of the ECM that occurs during development, axons reextended through pre-formed gaps that remained static over the three-day course of regeneration. Although axon guidance has traditionally been viewed as a product of chemical signaling, it is increasingly apparent that mechanical signaling in the form of stiffness gradients also regulates pathfinding (Koser et al., 2016). One possibility is that a gap in the ECM is sensed by the growth cone as a change in substrate stiffness and itself induces defasciculation. However, we have shown that the presence of an artificially made gap in the ECM is not enough to induce defasciculation of axons into the epidermis, and that axons prefer to grow through the original gap rather than the artificial one. This behavior implies the presence of a molecular cue that is necessary for attracting cut axons into the epidermis.

Our RNA *in situ* data confirmed the expression of *col18a1a* in hair cells as well as in the surrounding accessory cells of the neuromast. This pattern suggests that the hypothesized signaling complex would be distributed diffusely in the ECM and would only attract defasciculated growth cones into the general vicinity of the neuromast. Once inside the neuromast, growth cones must still distinguish hair cells from supporting cells, likely through contact with membrane-anchored synaptic markers on the basal surfaces of hair cells. Our bulk-sequencing data from denervated hair cells provide a good starting point for identifying candidate markers. Upon denervation, we observed upregulation of two genes that encode hair cell-specific synaptic adhesion molecules, *nrxn3b* and *lingo3a* (**Supplemental Fig. 8**). Neurexin3b has been shown to be essential for pairing pre- and post-synapses in neuromast hair cells (Jukic et al., 2024). Although Lingo3a is an uncharacterized orphan receptor, the Lingo protein family has been proposed to regulate nervous system development and axonal survival as do other leucine-rich repeat transmembrane proteins (Guillemain et al., 2020)..

In addition to restoring hair cell ribbon synapses with afferent fibers, there must also be recovery of modulatory efferent innervation. There are excitatory dopaminergic and inhibitory cholinergic inputs to neuromast hair cells by 5 dpf (Manuel et al., 2021; Toro et al., 2015). Dopaminergic axons do not make direct synaptic contacts with hair cells and are instead thought to release dopamine in a non-specific, paracrine fashion (Odstrcil et al., 2022; Toro et al., 2015). Cholinergic efferent terminals are in direct apposition to hair cells and suppress reafferent stimulation by signaling to hair cell nicotinic acetylcholine receptors containing α9 subunits (Odstrcil et al., 2022). Both dopaminergic and cholinergic efferents do not associate with the primordium, and instead follow the established path of towed afferents (Manuel et al., 2021; Toro et al., 2015). We assume that the regrowth of efferent fibers also likely depends on their association with regenerated afferent fibers, which greatly outnumber efferents in the nerve and in the neuromast (Dow et al., 2018). Once in the neuromast, different synaptic adhesion molecules on the hair cell must sort afferents to ribbon synapses and inhibitory efferents to post-synaptic membrane cisterns with acetylcholine receptors. Although afferent innervation is selective for hair cells of an identical orientation within a neuromast (Dow et al., 2018; Faucherre et al., 2009; Lozano-Ortega et al., 2018; Nagiel et al., 2008), efferent innervation is not, suggesting further differences in how the two synapses are reestablished. Agrin, another secreted HSPG that was upregulated in our bulk sequencing data, has a conserved role in clustering acetylcholine receptors and facilitating synaptogenesis at neuromuscular junctions through the Agrn/Lrp4/MuSK signaling pathway (Gribble et al., 2018; Kim et al., 2008; Zhang et al., 2008). While we did not see any gross defects in the afferent reinnervation of hair cells in *agrn* mutants, we cannot rule out impaired reinnervation by cholinergic efferents. Further work must be done to elucidate Agrin’s role in neuromast reinnervation, and more generally, the differences between efferent and afferent regeneration.

The simplicity and accessibility of the zebrafish posterior lateral line makes it an ideal model for studying the basic mechanisms of peripheral nerve regeneration and target selection. Here we present a model of one of the key steps of regeneration, the defasciculation of individual axons into the epidermis through gaps in the ECM. We identify *col18a1a* as an important gene for restricting axon branching to appropriate locations. A detailed understanding of the molecular interactions driving axon pathfinding and reformation of the sensory synapse has the potential to reveal therapies driving neural regeneration in otherwise permanently damaged sensory organs like the deafened cochlea.

## Acknowledgments

We thank Samantha Campbell for her expert care of the fish facility, Gaurav Shrestha for his assistance performing nerve lesions, Agnik Dasgupta for comments on axon-ECM interactions, and members of our research group for their comments on the manuscript. We also thank members of the Flow Cytometry Resource Center, Genomics Resource Center, and Bioinformatics Resource Center at The Rockefeller University for their support and guidance. The *Tg(HGn39d)* line was generated through the National BioResource Project (NBRP, Japan). R.S.R. was supported by a Medical Scientist Training Program grant from the National Institute of General Medical Sciences of the National Institutes of Health under award number T32GM152349 to the Weill Cornell/Rockefeller/Sloan Kettering Tri-Institutional MD-PhD Program and by a F30 Predoctoral Fellowship from the National Institute on Deafness and Other Communication Disorders of the National Institutes of Health under award number F30DC021350. A.J.H. is an investigator of Howard Hughes Medical Institute.

## Author Contributions

R.S.R conceptualized the project, conducted experiments, analyzed results, and wrote the initial draft of the manuscript. A.J.H. helped design experiments and interpret their results and edited the manuscript.

## Declaration of Interests

The authors declare no competing interests.

## Supplemental Information Titles

Supplemental File 1. Spreadsheet of differential gene expression in hair cells one day post nerve lesion

Supplemental File 2. Spreadsheet of differential gene expression in hair cells three days post nerve lesion

## Supplemental Figures and figure legends

**Supplemental Figure 1.**
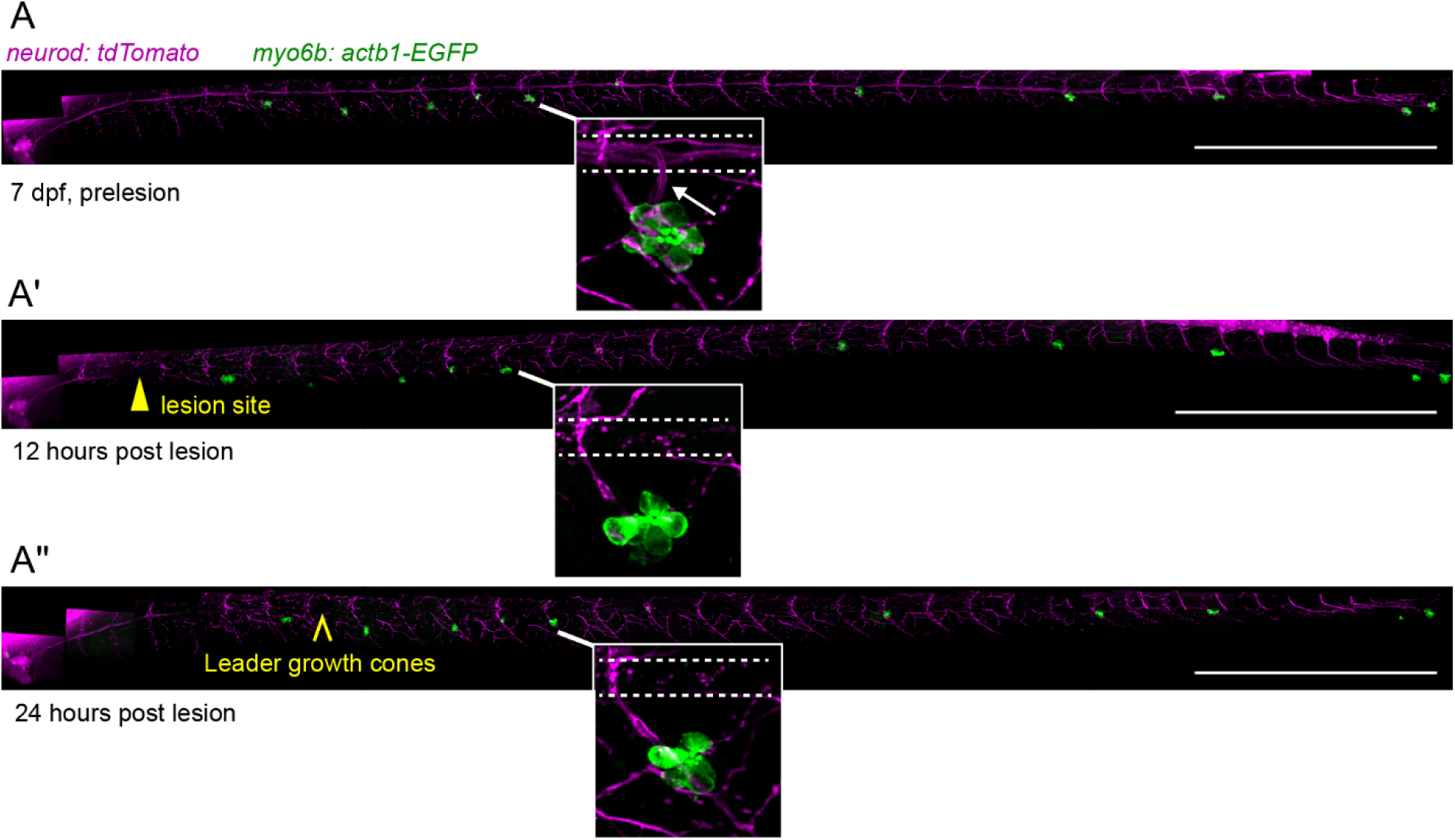
Hair cells post nerve lesion. (A) A representative 7 dpf *Tg(neurod:tdTomato, myo6b:actb1-EGFP)* larva before, 12 hpl (A’), and 24 hpl (A’’). *Neurod* is a pan-neuronal promoter driving expression of tdTomato (magenta) and the *myo6b* promoter selectively labels hair cells with GFP (green). At 12 hpl, cut axons had not yet traversed the lesion site (yellow arrowhead). At 24 hpl, leader growth cones (yellow caret) had partially extended along the horizontal myoseptum. The majority of pLL hair cells were distal to the cut axons and remained denervated (inset). Dotted white lines demarcate the pLL nerve and the white arrow denotes axon defasciculation prior to lesion. Scale bar, 500 µm.

**Supplemental Figure 2.**
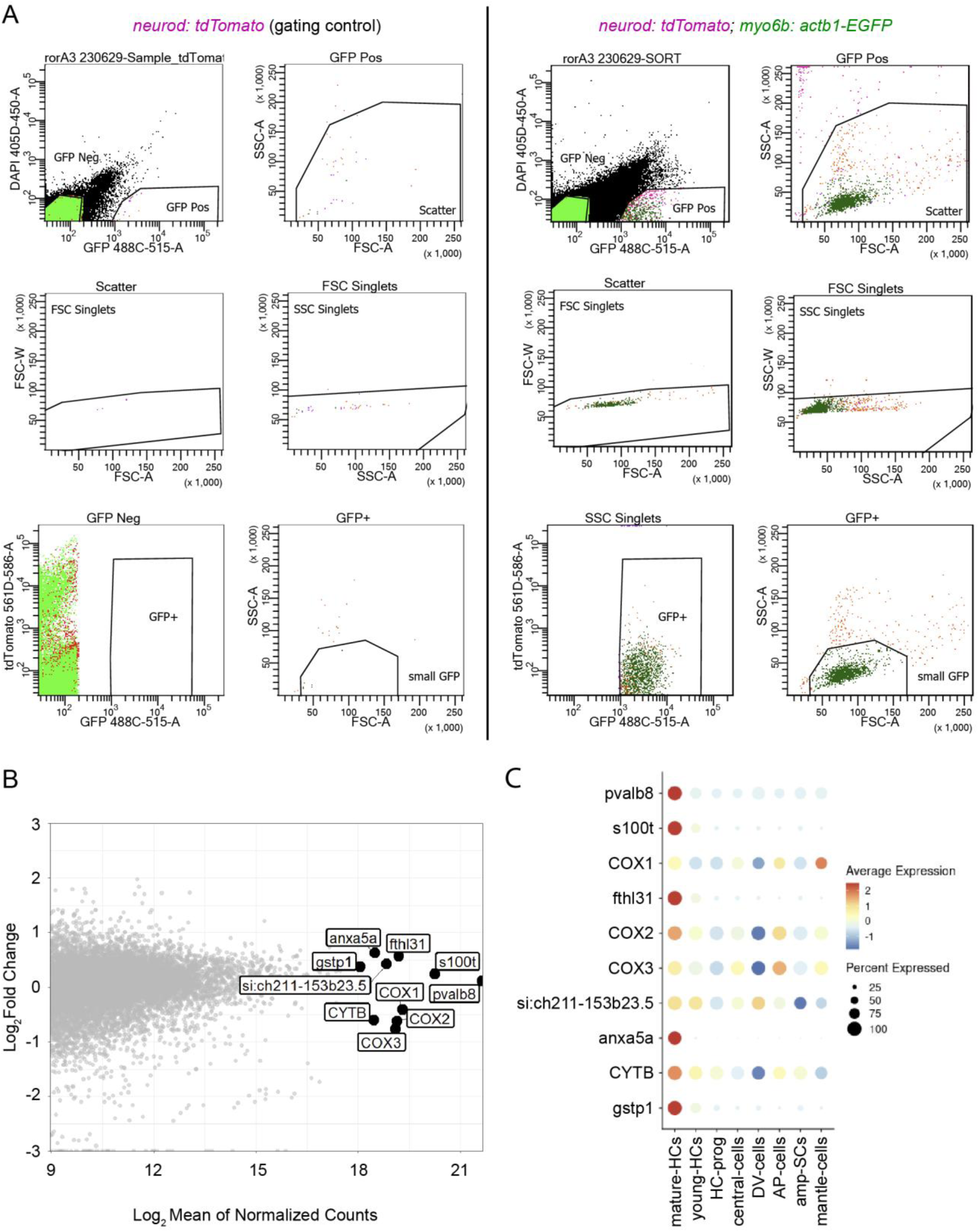
Isolation of hair cells for bulk sequencing. (A) The gating strategy used for fluorescence-activated cell sorting (FACS) of GFP-positive hair cells in *Tg(neurod:tdTomato, myo6b:actb1-EGFP)* larvae following nerve lesion (right). Singly transgenic *Tg(neurod:tdTomato)* larvae were used as gating controls for each session (left). FSC – forward scatter, SSC – side scatter. (B) An MA plot of transcripts sequenced in bulk hair cell samples 1 dpl compared to controls. The 10 most-expressed genes across all sample types are outlined in black. (C) The same genes in (B) cross-referenced to a published single-cell RNAseq dataset of the neuromast (Baek et al., 2022). The genes are selectively and highly expressed in hair cells, confirming the accuracy of the isolation and sorting protocol.

**Supplemental Figure 3.**
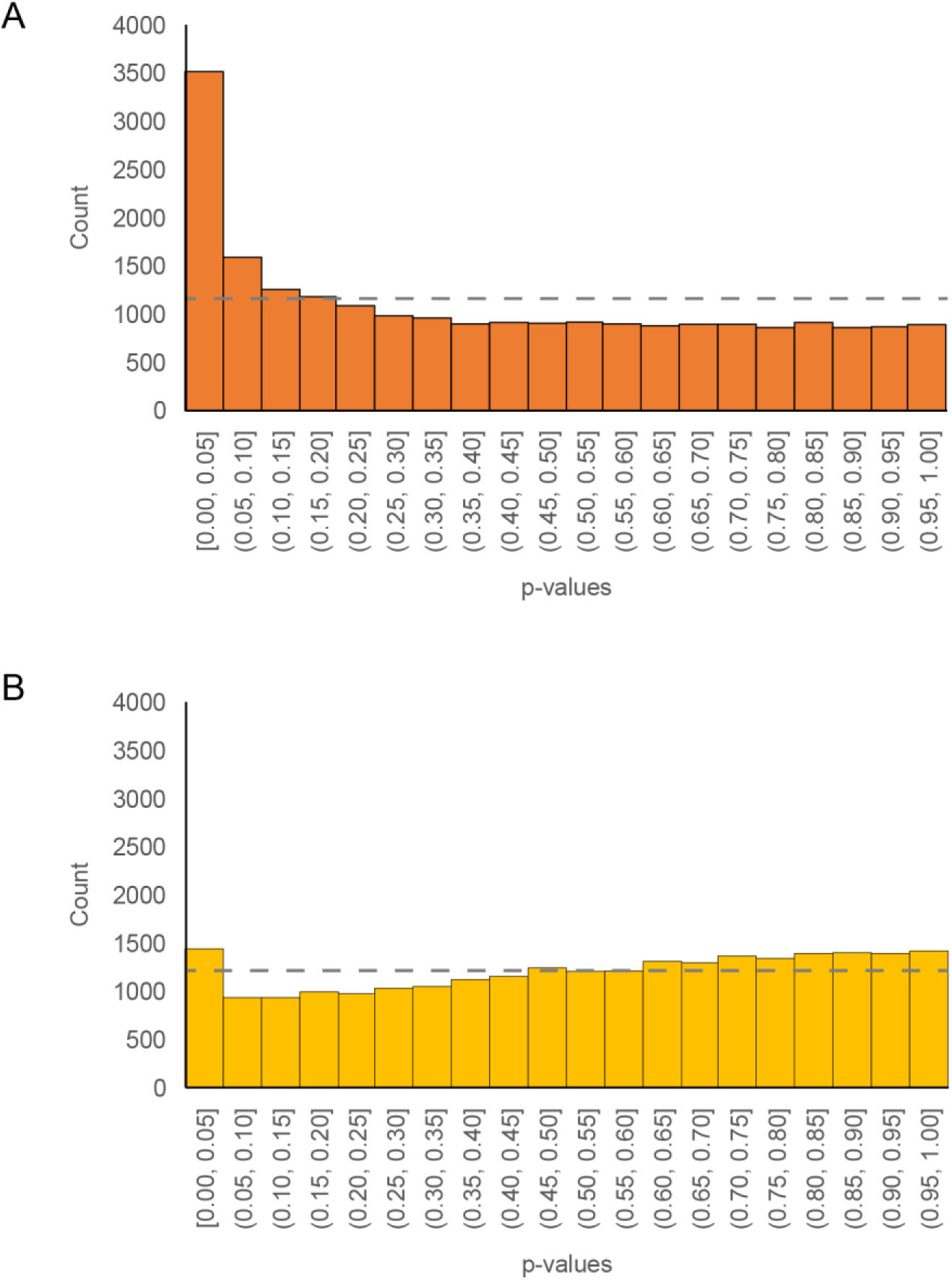
Distribution of p-values from bulk sequencing data. (A) The distribution of *p-values* from testing the differential expression of each gene in hair cells 1 dpl compared to controls. A left-sided peak of values below 0.05 deviates from a uniform distribution (gray dotted line). n = 22,148 total genes tested. (B) The distribution of *p-values* from testing the differential expression of each gene in hair cells 3 dpl compared to controls. The distribution more closely resembles a uniform distribution compared to (A). n = 24,209 total genes tested.

**Supplemental Figure 4.**
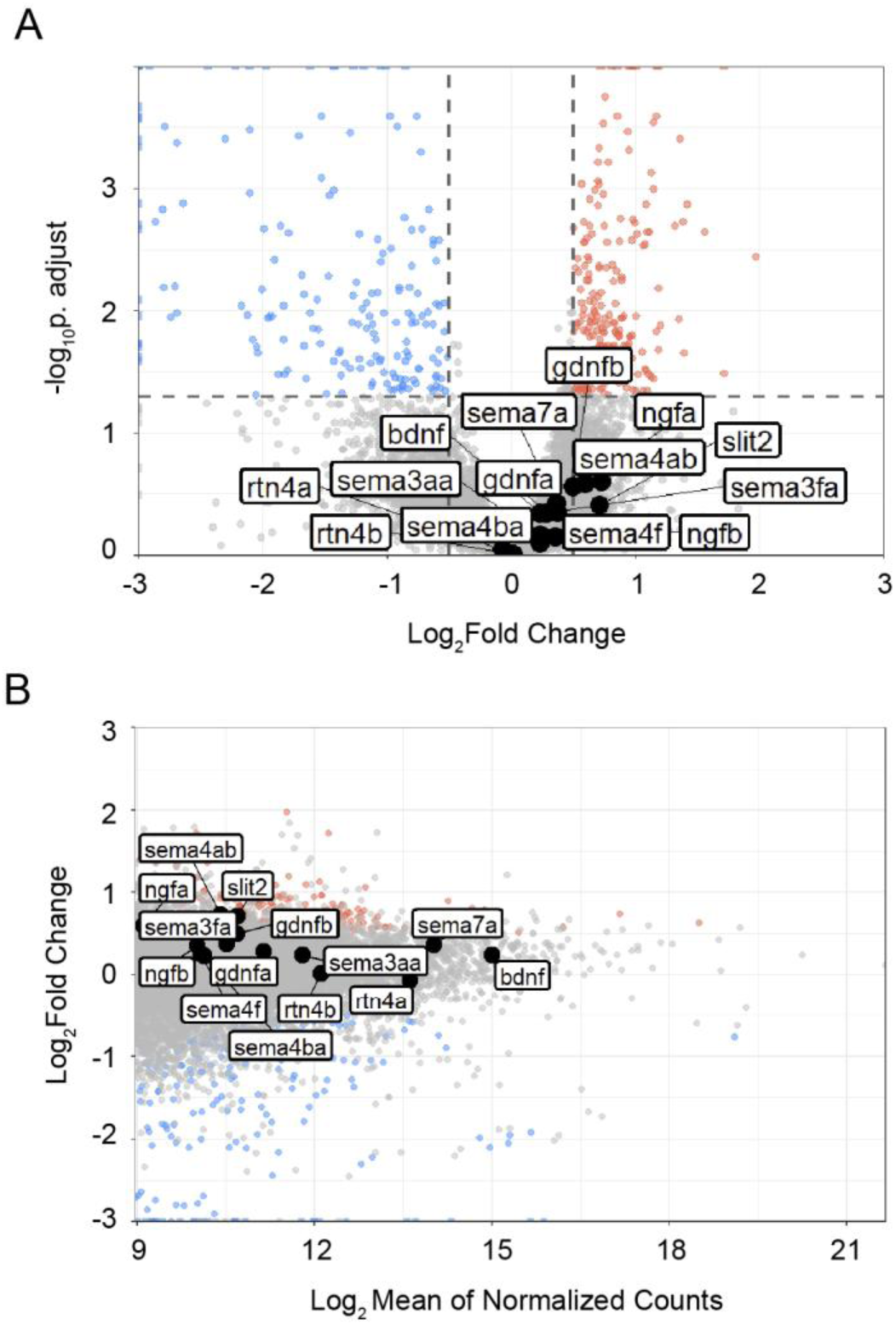
Expression of axon guidance cues in denervated hair cells. (A) A volcano plot showing canonical axon guidance cues are not differentially expressed in hair cells 1 dpl compared to innervated controls. Statistical significance set at a *p-value* of 0.05 adjusted for multiple comparisons (horizontal dashed line) and biological significance set at 0.5 log2fold change (vertical dashed line). (B) An MA plot of genes sequenced in hair cells 1 dpl compared to control with canonical axon guidance cues highlighted in black.

**Supplemental Figure 5.**
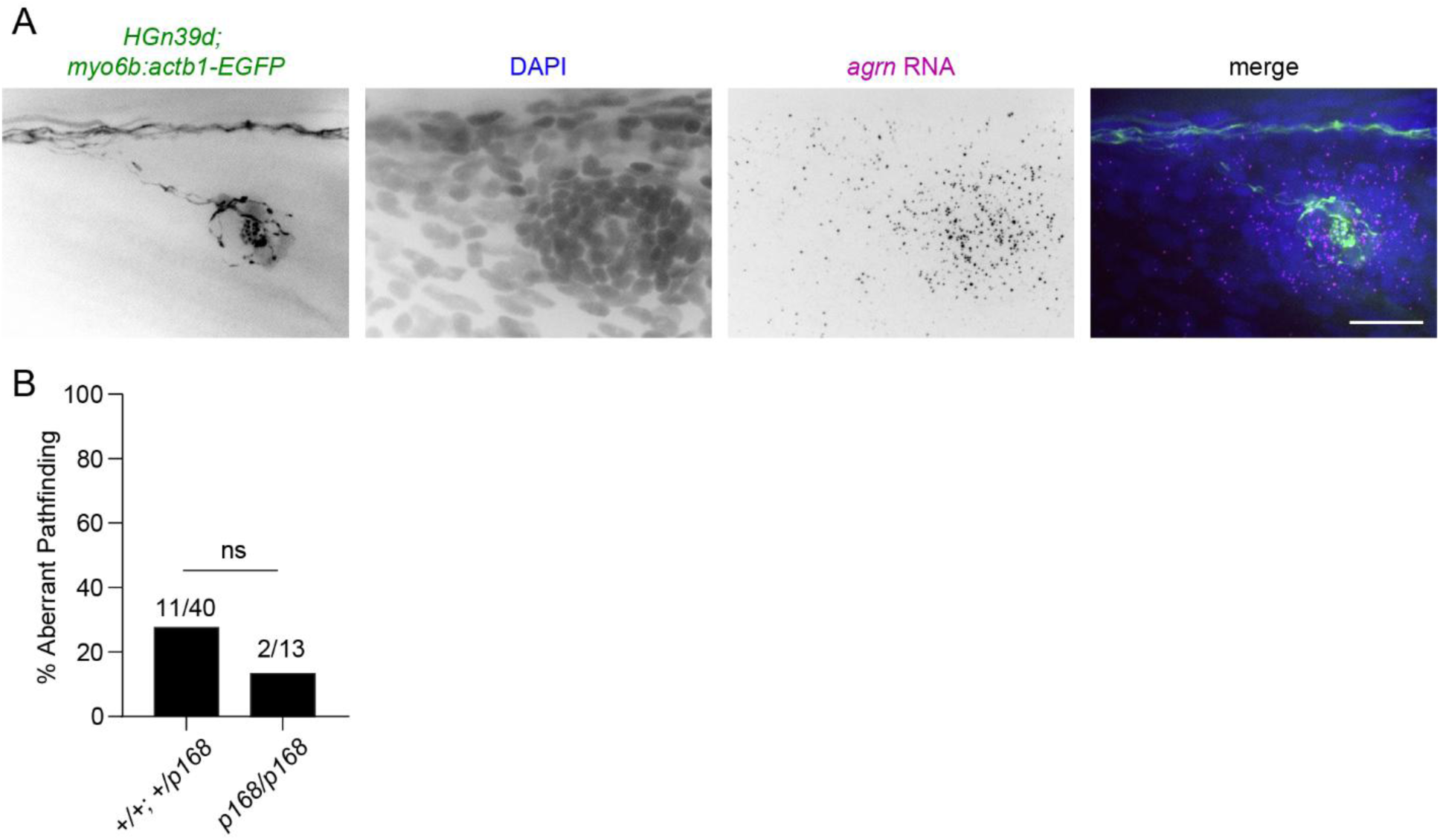
Normal nerve regeneration in *agrn^p168^* mutants. (A) RNA-FISH of *agrn* in a fixed 8 dpf *Tg(HGn39d; myo6b:actb1-EGFP)* larva. Expression of *agrn* is specific to hair cells and supporting cells of the neuromast. (B) There is an absence of aberrant afferent axon pathfinding during nerve regeneration in *agrn^p168/p168^ Tg(HGn39d; myo6b:actb1-EGFP)* mutants compared to siblings. n = 40 *+/+* and *+/p168* siblings, n = 13 *p168/p168* mutants. ns – not significant, Fisher’s exact test.

**Supplemental Figure 6.**
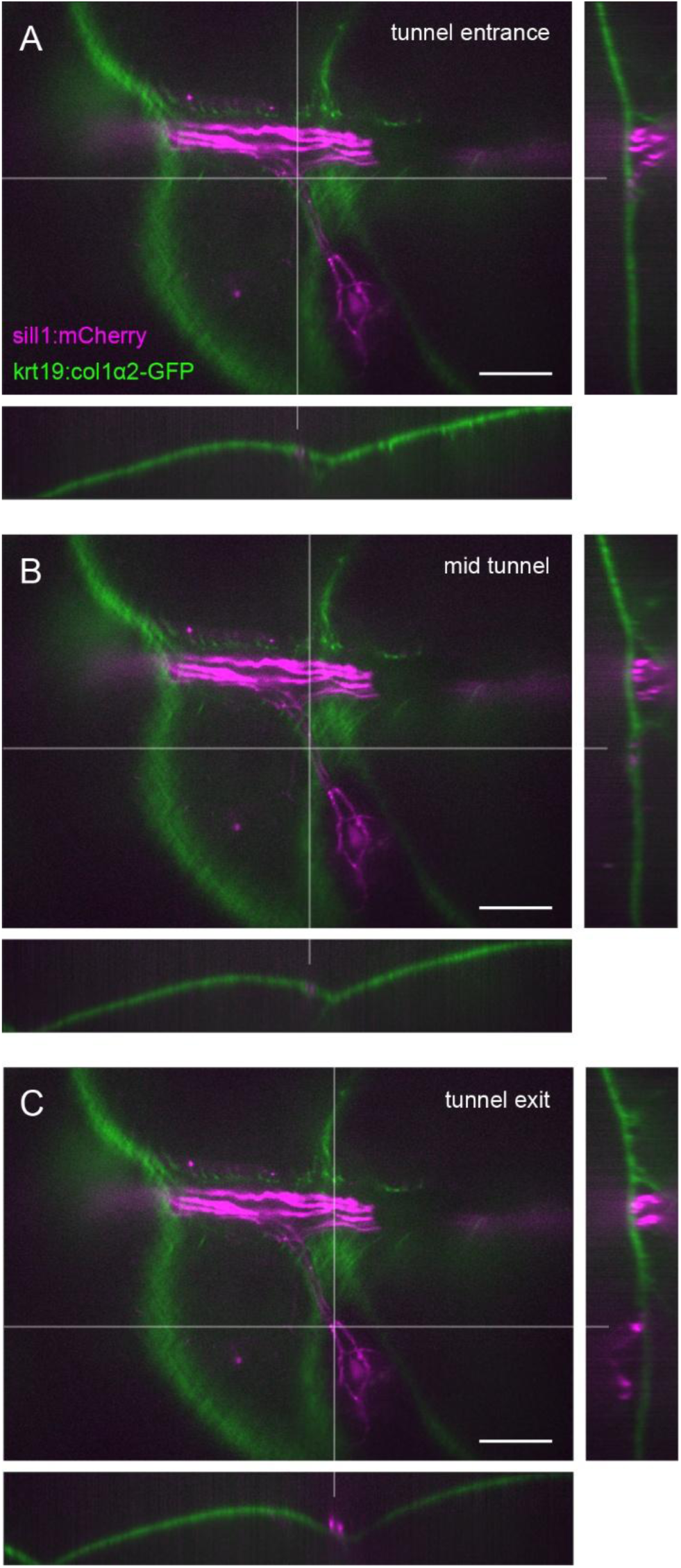
Innervation of secondary neuromasts through collagen tunnel. (A) Orthogonal views of the point where individual axons defasciculate from the axon bundle to innervate a secondary neuromast in a 5 dpf *Tg(krt19:col1α2-GFP, sill1:mCherry)* larva. The point of defasciculation coincides with the entrance to a tunnel within the collagen I matrix. Center – sagittal plane, right – axial plane, bottom – coronal plane. (B) Orthogonal views of the midpoint of the collagen I matrix tunnel. (C) Orthogonal views of the exit of the collagen I matrix tunnel at the secondary neuromast. All scale bars, 20 µm.

**Supplemental Figure 7.**
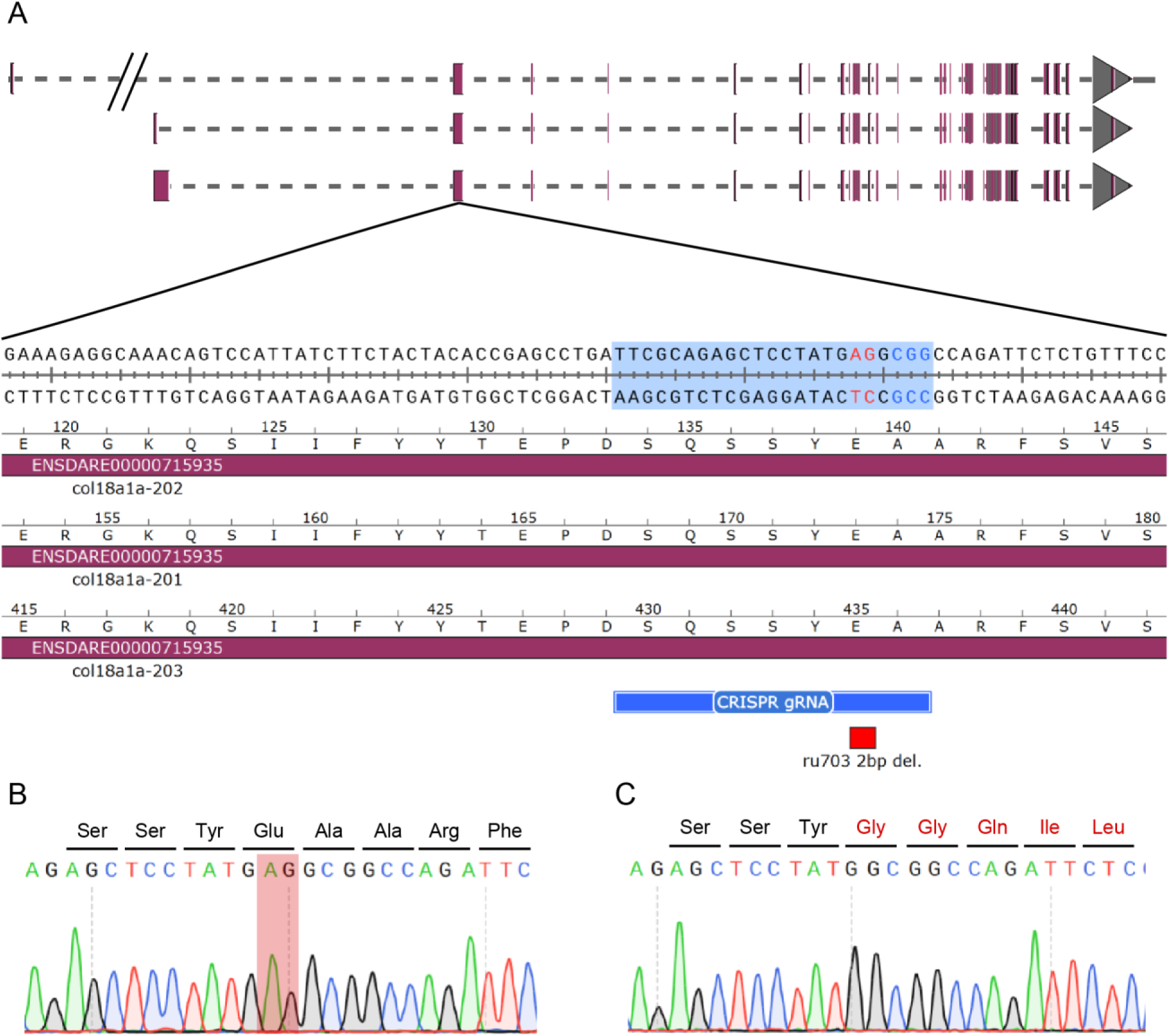
c*o*l18a1aru703 mutation. (A) The CRISPR gRNA sequence (highlighted) with adjacent CGG Cas9 PAM motif (blue) used to generate a stable *ru703* mutant line. The gRNA is complementary to either exon 2 in the 201 and 203 isoforms of *col18a1a*, and exon 3 in the 202 isoform. The two base pair AG deletion in *ru703* mutants are marked in red. (B) A chromatogram of a sequenced wildtype larvae. AG base pairs deleted in the mutated *ru703* allele are highlighted in red. (C) A chromatogram of a sequenced homozygous mutant *col18a1a^ru703/ru703^* larvae. The two base pair deletion creates a frameshift in the reading frame.

**Supplemental Figure 8.**
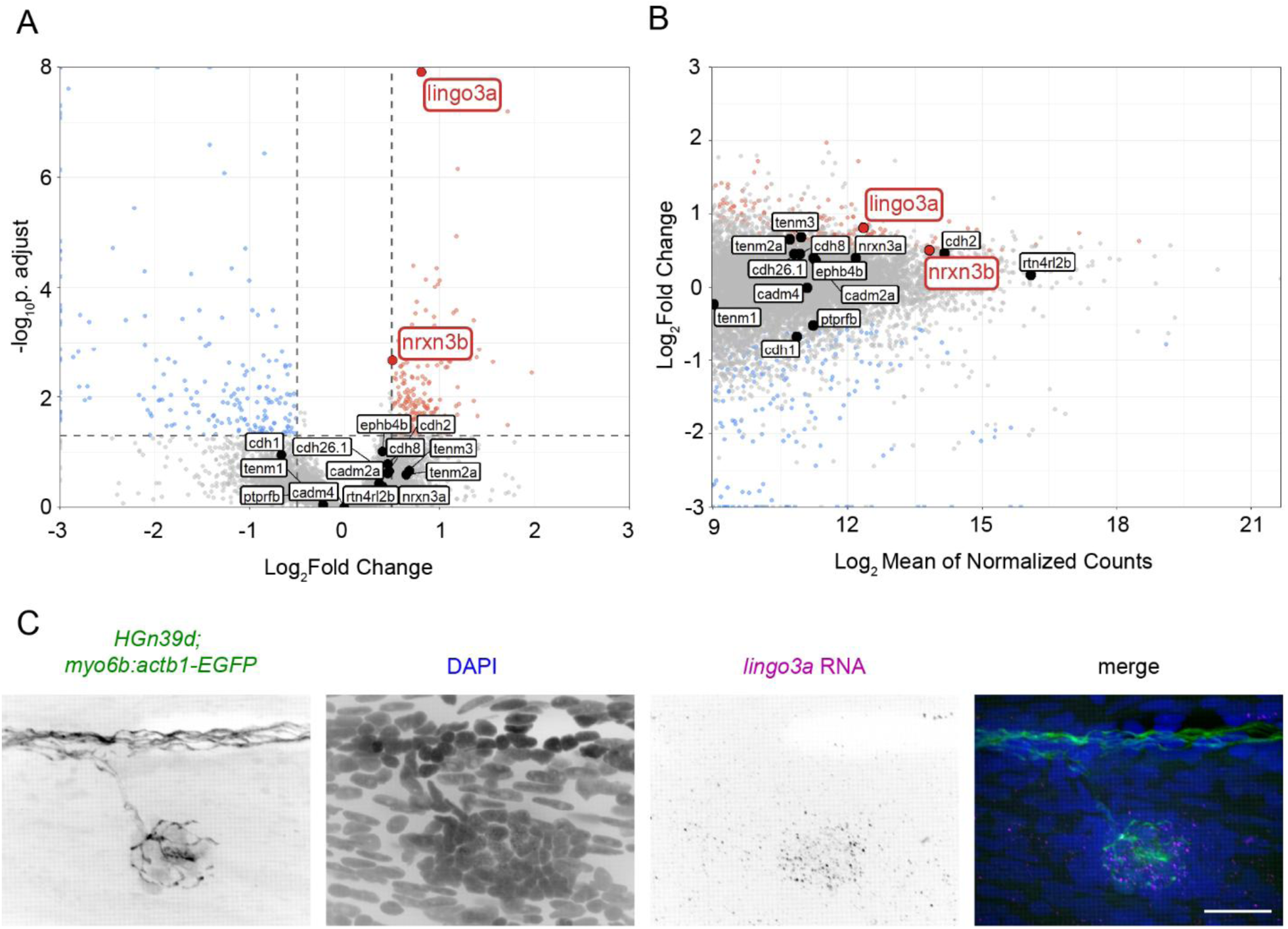
Synaptic adhesion genes upregulated in denervated hair cells. (A) A volcano plot with canonical synaptic adhesion molecule genes highlighted. *Lingo3a* and *nrxn3b* are upregulated (red) in denervated hair cells 1 dpl compared to innervated controls. Statistical significance set at a p-value of 0.05 adjusted for multiple comparisons (horizontal dashed line) and biological significance set at 0.05 log2fold change (vertical dashed line). (B) An MA plot of genes sequenced in hair cells 1 dpl compared to control with canonical synaptic adhesion molecule genes highlighted. (C) RNA-FISH against *lingo3a* in an 8 dpf *Tg(HGn39d; myo6b:actb1-EGFP)* larva reveals restricted expression in only hair cells of the neuromast. Scale bar, 20 µm.

## Materials and Methods

### Zebrafish strains

Zebrafish work was performed in accordance with animal protocol 22081-H reviewed and approved by The Rockefeller University’s Institutional Animal Care and Use Committee. Adult zebrafish (*Danio rerio*) were maintained under standard conditions (Nusslein-Volhard & Dahm, 2002). Naturally spawned and fertilized eggs were maintained in system water treated with 0.5 mg/L methylene blue (blue water) at 28.5°C on a 14-/10-hour light/dark cycle. Experiments were performed on 4-10 dpf larvae, before sex is determined.

The following previously described zebrafish transgenic and mutant lines were used: *Tg(cntnap2a)^nkhgn39dEt^* (Nagayoshi et al., 2008), *Tg(myo6b:actb1-EGFP)^vo8Tg^* (Kindt et al., 2012), *Tg(neurod:tdtomato)^vo12Tg^* (Toro et al., 2015), *Tg(krt19:colIα2-GFP)^zf2175Tg^* (Morris et al., 2018), *Tg(she:H2A-mCherry)^psi57Tg^* (Peloggia et al., 2021), and *agrn^p168^* (Gribble et al., 2018).

The *Tg(hsp70:NTR2.0-2A-mCherry en.Sill1)* line was created using the Gateway-based Tol2 kit (Kwan et al., 2007). Expression plasmids were created by combining *p5E- hsp70* (Kwan et al., 2007), *pME-NTR2.0-2A-mCherry* (Sharrock et al., 2022), *p3E-Sill1* (Pujol-Marti et al., 2012) and *pDestTol2pA2* (Kwan et al., 2007) plasmids. Verified plasmid DNA was injected at a concentration of 30 ng/µl into single-cell AB embryos along with 125 ng/uL of Tol2 transposase mRNA. The mutant *col18a1a^ru703^* line was created through CRISPR/Cas9 gene editing of wildtype embryos. Adults with a 2 base pair deletion in exon 2 (201, 203 transcript) or exon 3 (202 transcript) of the *col18a1a* gene (ENSDARE00000715935) were pooled and outcrossed to *Tg(HGn39d, myo6b:actb1- EGFP)* fish to generate heterozygotes identified by tail-clip genotyping with primers: *col18a1a^ru703^* forward 5’- GCGTCTCCAAAGTCTTCGAC-3’ and *col18a1a^ru703^* reverse 5’- GCGCTCGAAATTGACAACCT-3’. Similarly, *agrn^p168^* mutants were outcrossed to *Tg(HGn39d, myo6b:actb1-EGFP)* fish to generate heterozygotes identified by tail-clip genotyping with primers: *agrn^p168^* forward 5’- CAATGGTCAGAAGACAGACGG -3’ and *agrn^p168^* reverse 5’- GGCTCCACTGTATATTATGCTGC -3’.

### Lateral line nerve lesion and live imaging

Double transgenic *Tg(myo6b:actb1-EGFP; HGn39d)* larvae were anesthetized at 4 dpf in 600 µM tricaine (MS-222 tricaine methanesulfonate, Syndel) and embedded in 1% low melting point agarose for laser lesioning of the afferents of the right pLL nerve. All imaging was performed on a microlens-based, super-resolution confocal imaging system (VT-iSIM, Biovision Technologies) fixed to an Olympus IX81 frame and using a UPlanSApo 60x 1.3 NA silicone oil objective (Olympus). By convention, all displayed parasagittal images are oriented with the anterior of the fish to the left, the posterior of the fish to the right, the dorsal surface of the fish to the top, and the ventral surface of the fish to the bottom. All displayed coronal images are oriented with the anterior of the fish to the left, the posterior of the fish to the right, the lateral surface of the fish to the top, and the midline to the bottom. A 20 µm segment of the axon bundle at the 4^th^ somite was irradiated at three different confocal planes with an attached iLas Pulse laser system (Gataca Systems, France) equipped with a 355 nm laser, and complete transection of the bundle was confirmed 1 hour after ablation. The larvae were immersed in an imaging solution of 120 µM tricaine, 0.4 mg/mL pancuronium bromide (P1918, Sigma-Aldrich), and 1 mM sodium-L-ascorbate (11140, Sigma-Aldrich) diluted in system water for extended timelapse imaging of over 48 hours. Confocal stacks of the L1 neuromast were acquired at 1 µm intervals along the sagittal axis every 40 minutes.

The ErbB inhibitor AG1478 (658548, Sigma-Aldrich) was used to pharmacologically block Schwann cell signaling during axon bundle regeneration. The right pLL nerve was lesioned as described in 4 dpf *Tg(HGn39d)* larvae and the larvae were freed from the agarose and left to recover in either 1-4 µM AG1478 in 1% DMSO (Invitrogen; D12345) or only 1% DMSO in system water. Larvae were remounted for imaging 2 dpl, and the somite reached by the leading axon was recorded.

### Bulk RNA sequencing of denervated hair cells

The right and left pLL nerve was lesioned in 100-150 7 dpf *Tg(neurod:tdtomato; myo6b:actb1-GFP)* larvae. For sham lesions, tank siblings were exposed on both right and left sides to a UV LED for 30 seconds. Larvae were left to recover in blue water for either 1 or 3 days following lesion before tissue dissociation. We followed a modified neuromast tissue dissociation protocol (Baek et al., 2022) to generate a crude cell suspension for fluorescence-activated cell sorting (FACS). Larvae were anesthetized in 600 µM tricaine and decapitated with a fresh razor blade just anterior to the swim bladder to exclude hair cells from the anterior lateral line, otic vesicle, and dorsal branch of the pLL. The remaining larval trunks were rinsed with ice cold 1x DPBS (Gibco; 14190144)/0.04% BSA (03117332001, Roche) and transferred to a 15 ml polypropylene conical tube in 4.5 mL of ice-cold 0.25% trypsin-EDTA (Gibco; 25200056). The trunks were dissociated by trituration with a 1 ml pipette tip for 4 minutes on ice. The subsequent crude cell suspension was filtered through a 70-µm pore cell strainer into a 15 mL polypropylene round-bottom centrifuge tube. The cells were centrifuged at 2000 rpm for 5 min at 4°C. The supernatant was carefully removed from the cell pellet and the pellet was resuspended in ice cold 1x DPBS/0.04% BSA before re-centrifugation at 2000 rpm for 5 min at 4°C. This step was repeated once more, and the cell pellet resuspended in 500 µL of 1x DPBS/0.04% BSA. This suspension was filtered through another 70-µm pore filter into a 1.5 mL screwcap tube with 100 units of Protector RNase Inhibitor (3335399001, Roche). The total time between larvae decapitation and cell sorting was kept to under 90 minutes to minimize hair cell death.

GFP-positive cells were sorted at the Rockefeller University Flow Cytometry Resource Center with a BD FACSAria II (BD Biosciences) using a 100 µm nozzle at 20 psi. The 405, 488, and 561 nm laser excitation lines with 450/40 nm, 515/20 nm, and 586/15 nm emission filters respectively were used for collection. For every sorting session, a fresh control sample of GFP-negative cells was used to establish gating parameters. GFP-positive hair cells were sorted based on size and GFP fluorescence directly into RLT+ cell lysis buffer (QIAGEN) with 1% beta-mercaptoethanol. Total sorting time was limited to 30 minutes to limit hair cell death. Total RNA was extracted after sorting using the RNeasy Plus Micro Kit (QIAGEN) and stored at -80 °C until sequencing. Hair cell RNA from three technical replicates were collected for each condition: 1 day post nerve lesion, 1 day post sham lesion, 3 days post nerve lesion, 3 days post sham lesion. The yield and quality of isolated RNA was measured with a bioanalyzer (Agilent 2100). All samples had an RNA integrity score (RIN) greater than 8.0 and were subsequently sequenced by the Rockefeller University Genomics Resource Center. 1 ng of total RNA was used to generate full length cDNA using the Clontech Smart-Seq v4 Ultra Low Input RNA kit (634888, Clontech). 1 ng of cDNA was then used to prepare libraries using the Illumina Nextera XT DNA sample preparation kit (FC-131-1024, Illumina). Libraries generated from hair cells 1 day post nerve lesion and control hair cell samples 1 day post sham lesion were sequenced together on an Illumina NextSeq500 with single end 75-base pair reads. Libraries generated from hair cells 3 days post nerve lesion and control hair cell samples 3 days post sham lesion were sequenced together on an Illumina NextSeq 2000 with single end 100-base pair reads. All samples were sequenced to a depth of approximately 66 million reads.

Quality control and processing of the raw bulk RNA sequencing data was performed by the Rockefeller Bioinformatics Resource Center using a standardized bulk sequencing analysis pipeline programmed in R. Briefly, reads were aligned to the zebrafish reference genome (GRCz11) using the Rsubread package and transcript counts quantified with Salmon and the tximport package. Quality control metrics were assessed with the picard package. Processed data was normalized and statistical testing was performed using the DESeq2 package. Only genes with normalized transcript counts above 500 were included in volcano plots and MA plots. Gene set enrichment analysis was performed using the fgsea package (Korotkevich et al., 2021). All raw sequencing data has been deposited in the Gene Expression Omnibus (GEO) database (Accession: GSE297706). Detailed step-by-step workflows, quality control metrics, and additional R code used for the processing of bulk sequencing data in this study are publicly available (https://github.com/rsroy27/CollagenXVIII_2025_Manuscript). Gene lists and normalized transcript counts for each sample is included as an Excel table in the supplementary material of this paper.

### RNA Fluorescence *in situ* hybridization

RNA fluorescence *in situ* hybridization was performed on whole larvae using standardized hybridized chain reaction (HCR) v3 and v4 protocols, reagents, and probes (Molecular Instruments). Larvae were fixed in 4% paraformaldehyde for 24 hours at 4°C either one day post nerve lesion or with the nerve left intact. Fixed larvae were washed in PBS and dehydrated in 100% methanol before storage at -20°C overnight. Larvae were then rehydrated in graded methanol/0.1% PBS-Tween 20 washes, treated with 10 µg/mL proteinase K for 10 minutes, and postfixed with 4% paraformaldehyde for 20 minutes. Double transgenic *Tg(HGn39d; myo6b:actb1-EGFP)* 8 dpf larvae were simultaneously probed for *col18a1a* (NM_001349195.1) and *sox10* (NM_131875.1) mRNA either one day post nerve lesion or in larvae where the nerve was left intact. Fixed *Tg(HGn39d; myo6b:actb1-EGFP)* larvae were also separately probed for *agrn* (NM_001177452.1) and for *lingo3a* (NM_001082854.1) mRNA. Single transgenic *Tg(krt19:col1α2-GFP)* 8 dpf larvae were probed for *col18a1a*. The proteinase K incubation step was skipped for these larvae to better preserve the structure of the ECM. Larvae were incubated with RNA probes overnight at 37°C followed by incubation with corresponding fluorescent hairpin markers overnight at room temperature. To aid in cell counting, a 1:200 dilution of 1 mg/mL DAPI was used in the final wash step before mounting larval trunks in ProLong Diamond Antifade (Invitrogen; P36961) under a 1.5 coverslip. Confocal stacks of individual neuromasts were obtained at 0.200 µm intervals along the sagittal axis. Contrast adjustments, cell counting, and three-dimensional distance measurements were performed in Fiji and Imaris.

### Extracellular matrix imaging and puncture

The collagen I epidermal boundary layer in relation to pLL nerve afferent axons was imaged in live 5 dpf *Tg(krt19:col1α2-GFP; hsp70:NTR2.0-2A-mCherry en.Sill1)* larvae. The collagen I epidermal boundary layer in relation to primordium-derived neuromast and interneuromast cells was imaged in live 4 dpf *Tg(krt19:col1α2-GFP; she:H2A-mCherry)* larvae. All confocal stacks were obtained at 0.200 µm intervals along the sagittal axis. Gap area measurements, three-dimensional distance measurements, and orthogonal slicing of the confocal stack was performed in Fiji and Imaris.

Confocal stacks of primary L3 or L4 neuromasts were obtained in *Tg(krt19:col1α2- GFP; hsp70:NTR2.0-2A-mCherry en.Sill1)* larvae pre-nerve lesion at 5 dpf, 1 day post lesion at 6 dpf, and 3 days post lesion at 8 dpf. In between image acquisition, larvae were freed from the agarose used for immobilization and left to recover in blue water. The somite number at which a neuromast was imaged pre-nerve lesion was used as a fiduciary marker for repeat imaging of the same neuromast 1 day and 3 days post lesion. Stacks were viewed in Imaris to assess the number of neuromasts reinnervated through the pre-existing gap in the ECM.

A glass microneedle pulled to a diameter of approximately 20 microns was mounted on a stereotactic rig to precisely puncture the skin on the lateral side of 5 dpf *Tg(krt19:col1α2-GFP, hsp70:NTR2.0-2A-mCherry en.Sill1)* larvae somewhere along the horizontal myoseptum, directly following nerve lesion. Larvae were then left to recover in blue water for 3 days before imaging. The artificial gap created by needle puncture was located by scanning along the length of the regenerated pLL nerve for a large, jagged- edged hole in the collagen I matrix. Because the larvae were poked at a random location along the horizontal myoseptum, the artificially made gap occurred either on an open region of the nerve with no neuromasts nearby or on a region of the nerve close to a neuromast. In the latter case, the artificial gap was either anterior to, posterior to, or encompassed the natural, pre-existing gap in the ECM. Larvae in which punctures coincided directly with the natural gap were excluded from analysis. Confocal stacks were viewed in Imaris to assess the extension of axons into the epidermis through artificial gaps. Epidermal extension was defined as complete passage of an axon into the epidermis, and growth beyond the borders of the gap.

### Mutant generation and phenotype scoring

The *col18a1a^ru703^* line was created through CRISPR/Cas9 gene editing of wildtype embryos and selection of a stable germline mutation following standard protocols (Kroll et al., 2021). Potential target sites for gene editing were determined using ChopChop (v3 https://chopchop.cbu.uib.no/). A 14.25 µM ribonucleoprotein complex of sgRNA (IDT) designed against an exon common to all three major isoforms of *col18a1a* and high fidelity Cas9 (IDT; 1081061) was injected into the yolk of single-cell embryos. Adult founders, with germline transmission of a 2 bp deletion resulting in a frameshift and premature stop codon in exon 2 (201, 203 isoform) or exon 3 (202 isoform), were identified by Sanger sequencing of offspring and use of the PolyPeak Parser tool (Hill et al., 2014). These adult founders were outcrossed to *Tg(HGn39d; myo6b:actb1-GFP)* adults and F1 heterozygote offspring were raised to sexual maturity. All experiments were performed on F2 larvae with fluorescent lateral line afferent neuron and hair cell labels. Similarly, *agrn^p168^* mutants with a 7 bp deletion/2 bp insertion in the donor splice site of exon 31 (Gribble et al., 2018) were outcrossed to *Tg(HGn39d, myo6b:actb1-GFP)* adults to incorporate a lateral line afferent and hair cell marker. Heterozygotes were raised to sexual maturity and experiments performed on offspring.

Axon bundle width was measured on maximum projections of confocal stacks taken from *col18a1a* mutants and siblings at the level of the 10^th^ and 15^th^ somite. We did not observe any phenotypic differences between *agrn^+/p168^* and *agrn^+/+^* siblings, or *col18a1a^ru703/+^* and *col18a1a^+/+^* siblings, and thus grouped them together for analysis. For both *agrn^p168^* and *col18a1a^ru703^* mutant larvae and wildtype siblings, the right pLL nerve was lesioned at 5 dpf and larvae left to recover for 3 days in blue water. At 8 dpf, the regenerated pLL nerve was scanned for aberrant axon pathfinding. Aberrant pathfinding was defined as the inappropriate defasciculation of axons at locations where there were no neuromasts or the extension of axons beyond the terminal neuromast. The observer was blinded to the genotype of the larvae, which was determined through Sanger sequencing of larvae after phenotype scoring. Each experiment was repeated on at least three separate clutches.

### Statistical Analysis

Statistical testing for differential gene expression in the bulk RNA sequencing data was performed using the DESeq2 package in R, employing the Wald test with a correction for multiple comparisons using the Benjamini and Hochberg method. Normalized enrichment scores were generated using the fgsea package in R, using the fast gene set enrichment analysis method with a correction for multiple comparisons using the Benjamini and Hochberg method. Additional analysis was performed in GraphPad Prism 10 using Student’s t-test for comparison of quantitative data, Chi-Squared test for testing independence between categorical variables, and Fisher’s exact test when sample size assumptions for the Chi-Squared test were not met. Statistical details such as sample size and significance cutoffs are given in the respective Figure legends.

### Data and Resource Availability

All unique plasmids, transgenic fish lines, and mutant fish lines generated in this study are available from the corresponding author without restriction. Due to size limitations, all raw experimental image data analyzed in this paper will be shared by the corresponding author upon request. Quality control and analysis of bulk RNA sequencing data was performed with custom code written by The Rockefeller University Bioinformatics Resource Center and is publicly available on Github (https://github.com/rsroy27/CollagenXVIII_2025_Manuscript). Bulk RNA-seq data have been deposited in the Gene Expression Omnibus (GEO) (Accession: GSE297706) and is publicly available.

## Notes

https://github.com/rsroy27/CollagenXVIII_2025_Manuscript

https://www.ncbi.nlm.nih.gov/geo/query/acc.cgi?acc=GSE297706

### Competing Interest Statement

The authors have declared no competing interest.

